# Quantifying the impact of groundwater fauna and temperature on the ecosystem service of microbial carbon degradation

**DOI:** 10.1101/2025.07.26.665685

**Authors:** Susanne I. Schmidt, Nora Rütz, Jürgen Marxsen

## Abstract

Groundwater ecosystems fulfil functions that humankind relies upon, e.g. for sustainable drinking water production. Quantification of these services is lacking so far. Thus, it is not possible to predict scenarios (e.g. future climates). Based on data from a comprehensive groundwater ecosystem study comprising four zones of varying land use and groundwater / surface water exchange, we parameterized a quantitative dynamic food web model (recharged organic carbon, microorganisms using the biodegradable fraction of this carbon, and fauna grazing on the microorganisms). With the model satisfactorily reflecting the field data, we calculated five further scenarios, three of which without fauna (mortality e.g. due to contamination, sudden peaks of temperature etc.). Two of the “fauna” and two of the “no fauna” scenarios were run with temperature elevated by 1.5°C and 3°C, respectively. The ecosystem service of carbon degradation was expressed as the difference in carbon concentration between the beginning of the simulation and the end of the simulation. In most scenarios, remaining carbon increased over time. The remaining carbon in some scenarios was up to 6.6 times as high in the “no fauna” scenarios compared to the reference case. Fauna was thus shown to fulfil a service by promoting microbial carbon degradation that may be substantial. Sustainable drinking water production is more reliable and less costly, the more active the groundwater fauna in the production area is. This model set up can serve to test other cases of varying physical and chemical variations and disturbances.

## 1 Introduction

Ecosystems fulfil functions that humans rely on. Groundwater is no exception (Avramov et al., 2010). The food web in groundwater (Malard et al., 2023) is often truncated (Gibert and Deharveng, 2002), but may encompass several trophic levels (Ercoli et al., 2019; Malard et al., 2023), depending on the respective environmental conditions. One of the functions that these food webs perform is processing organic carbon. This function of self-cleaning in groundwater, i.e. degrading carbon compounds to ultimately the gases CO_2_ or CH_4_, is probably most relevant to humans. These functions are one prerequisite for sustainable drinking water production, which is one of the 17 goals (goal 6) of the 2030 Agenda for Sustainable Development (United Nations General Assembly, 2017).

Some groundwater food webs are qualitatively well described, in terms of e.g. “who eats whom” (Bradford, 2009; Bradford et al., 2010; Ercoli et al., 2019; Francois et al., 2016; Leijs et al., 2015). Quantitative evaluations are a lot scarcer (but see e.g. Foulquier et al., 2011). To this date, no quantitative evaluation of ecosystem services in groundwater and their susceptibility to temperature existed. However, we urgently need to quantify the impacts of temperature on groundwater ecosystem services since we may have to future-proof drinking water production if groundwater ecosystems change their functionality due to increasing temperatures and changing climates and land uses.

Generally, at the bottom of a food web are producers. In groundwater, where light is lacking, however, primary production is only possible in the form of chemolithoautotrophy. While this is more prevalent than previously thought (Kellermann, 2008), heterotrophy, i.e. using organic resources that enter the groundwater ecosystem with groundwater recharge, will dominate in most places (Herrmann et al., 2015). Where heterotrophic degradation of imported organic material yields sufficient energy, a further level of the food web may be supported, i.e. grazers consuming producers (we use the term producers here to encompass autotrophs and heterotrophs feeding on imported organics). Where even the producer biomass becomes substantial, a further level may be added, i.e. predators preying on grazers (Ercoli et al., 2019; Saccò et al., 2019). There are few taxa with just one feeding mode – most fauna will be grazing for at least part of their life cycle (Hose et al., 2022). It can be assumed that the biomass and energy in a respective level reduces to 10 % of the underlying level’s biomass and energy (Lindeman, 1942; Odum, 1957). The loss is due to respiration which occurs due to mobility, reproduction, hunting etc. (Kozłowski, 1992), and in a full budget, would result in outgassing of CO_2_ (or CH_4_ in an anoxic system; Herrmann and Taubert, 2022; Nydahl et al., 2020) from the respective compartment, as a result of respiration (Moinet et al., 2023). Under which circumstances which complexity of the food web emerges, is still largely unknown and has thus been near-impossible to parameterize.

It is well-known that grazing either reduces (Kota et al., 1999) or advances (Mattison et al., 2005) bacterial biodegradation of contaminants. Thus, adding levels to the food web might increase or it might compromise remediation and attenuation.

While structuring food webs with just producers, grazers, and predators is grossly simplifying, this pattern has successfully been used e.g. in conceptualizing biomanipulation where e.g. predatory, piscivorous fish are added to a water body which harbours producing algae, grazing zooplankton and zooplankton-consuming planktivorous fish (Carpenter et al., 1985; Vrba et al., 2024). In such systems, commonly, the planktivorous fish reduce the zooplankton so that the algae are not grazed upon, may grow to harmful algal blooms and thus, deteriorate the water quality (Dadi et al., 2023). By adding piscivorous fish, the planktivorous fish are reduced, thus, more zooplankton remains which grazes more efficiently the algal blooms (Triest et al., 2016). Adding piscivorous fish is thus one measure for improving water quality. Is such management possible in groundwater as well? Or, vice versa, will the loss of grazing fauna result in lower water quality?

In drinking water production, it is advantageous to have raw water with low organic substances so that little respiration (which would reduce the dissolved oxygen) takes place in the pipes on the way to the customers. Where dissolved oxygen is low or lacking, respiration of organic substances may lead to CH_4_ which may be harmful to organisms and might thus limit degradation further. Anoxic conditions, resulting from respiration of excessive carbon sources, may lead to reduction of substances, and thus, dissolution of compounds which are bound in macro molecules at oxic conditions. E.g. aluminium, iron, etc. may become soluble at anoxic conditions (Dadi et al., 2023), and may be toxic to organisms. Thus, high raw water quality, with low levels of carbon, begins in the aquifers with a community of active organisms, degrading the recharged carbon and ensuring high oxic state throughout the drinking water production.

In addition to reduced conditions leading to undesired dissolved compounds, recently, brownification has become increasingly a problem (de Wit et al., 2016). One source of high molar mass organic compounds colouring the water yellow or brown is forest die back (Kopáček et al., 2023). The combination of dry summers stressing trees, and bark beetles making use of higher temperatures to produce more offspring than at colder climates (Seidl and Rammer, 2017) have led to expansive forest die back, e.g. in the Hercynian mountains (Musolff et al., 2024) in Germany and the Bavarian and Šumava mountains at the German-Czech boundary (Schmidt et al., 2024). This sudden increase of biomass is broken down to organic dissolved compounds many of which are undesired in drinking water production.

But higher temperatures have more, and more direct, effects than just favouring bark beetle generation times and dry summers. Dissolved oxygen solubility in water decreases with increasing temperatures (Retter et al., 2020), and brownification is believed to be exacerbated by increased temperatures (Blanchet et al., 2022). For these, and further, reasons, the temperature regime in a groundwater body is important when evaluating the current and potential future state of an aquifer. Globally, air temperature has increased by 1.5 °C compared to pre-industrial times in 2025 (Bevacqua et al., 2025). The temperature increase in groundwater is delayed (Benz et al., 2017) and depends strongly on soil and geological properties and hydrological processes (Rau et al., 2015), but has, e.g. in Austria, reached approximately 0.4 ± 0.5 K per 10 years, meaning that an increase of 1.5 °C might be observable within 40 years. Benz et al. (2024) predicted that between 2000 and 2099, groundwater will warm on average by 2.1°C, under a medium emission pathway. It is thus important to estimate which effect these temperature increases will have on the groundwater ecosystem and the functions it fulfils.

On top of increased carbon input and higher likelihood for reduced conditions, increasing temperatures also impact groundwater organisms directly. E.g., increasing temperatures were related to increased microbial biomass in a temperature plume in Southern Germany, with the microbial community changing qualitatively (Brielmann et al., 2009). A change in microbial ecosystem function might follow but has so far rarely been characterized.

The microbial community is main food source for fauna. Groundwater fauna are less sensitive to some contaminants such as heavy metals than their surface relatives (Mösslacher, 2000), but are very sensitive towards temperature. Faunal diversities decreased with increasing temperatures within a temperature plume (Brielmann et al., 2009). This has implications on ecosystem services. Avramov et al. (2013) demonstrated the response of crustacean stress hormones to increasing temperatures. Groundwater niphargids and asellids differed in the temperature range they tolerated, with niphargids covering a wider range (Brielmann et al., 2011). Spengler and Hahn (2018) identified a threshold value for middle European groundwater systems of 13 °C above which fauna community changed drastically in composition and presumably function. Avramov (2013) and Brielmann et al. (2011) performed tests analogous to toxicity tests and extrapolated lethal temperatures for various durations of exposure. Like for the favoured temperature ranges, niphargids survived longer at higher temperatures than asellids. The lethal temperatures from these experiments can be used to parameterize fauna temperature-related mortality in models. Effects of temperature on the sublethal level, e.g. dissolved oxygen consumption, have also been quantified (Di Lorenzo & Galassi, 2017) and should be included in future models – here, we concentrate on the lethal temperature effect on fauna only.

In extreme cases, in summary, a temperature increase of a few °C may be lethal to groundwater fauna. Groundwater fauna is thus threatened from extinction, like, globally, a quarter of freshwater fauna (Sayer et al., 2025). In contrast to climate opportunists following along the preferred temperature range (Dormann et al., 2018), groundwater fauna’s expansion or retreat is limited by the aquifers’ physical boundaries. For the same reason, any faunal community not coping with increased temperatures will probably not be replaced by a higher-temperature-adapted community. With fauna’s ecosystem function lacking, any effect of microorganisms being more active at higher temperatures, and thus degrading more carbon, may be (more than) compromised by the lack of grazing and thus rejuvenating microbial growth (Mattison et al., 2005).

Since it is hard to derive the relevant data sets on the required temporal and spatial resolution, while controlling for different environmental settings globally, one way to approach answers to these questions are models. Here, we present a model based on data from one of the most comprehensive data sets on a groundwater ecosystem (Husmann and Marxsen, 1988; Marxsen et al., 2021). Marxsen et al. (2021) grouped the wells in the River Fulda floodplain into four groups: one group combining wells near the river Fulda and in intense exchange with the river (“R”), one group covering wells that were characterized by a plume of a carbon source from the near-by village (“P”), one group characterized by agriculture with elevated nitrogen and phosphorus concentrations (“A”), and a group representing wells of a mixing zone being situated in-between the other three zones and combining characteristics of all zones (“M”). While many groundwater taxa are ubiquitous, neighbouring aquifer zones may differ markedly in fauna taxa composition (Di Lorenzo, Amalfitano, et al., 2025). The River Fulda data set thus enables to study reactions to scenarios of increased temperatures and / or setting fauna to 0 as if after a calamity, in different environmental exchange conditions, under the same climate. The data set allowed to check whether the developed model represented a range of measured field conditions. Once satisfied with the model, it is possible to change parameters and / or exclude fauna biomass to run scenarios representing warmer climates, and comparing ecosystem functions (here: degradation of organic carbon) when fauna is present versus absent. Fauna being absent might also be a consequence of climate change, since, as described, fauna does not survive well elevated temperatures.

The hypotheses were: 1) Without fauna, a groundwater ecosystem shows lower carbon degradation, thus, higher remaining organic carbon. 2) An increase in temperature by 1.5°C or 3°C leads to an increase in microbial degradation, and thus, lower remaining organic carbon. 3) Fauna is compromised more by increasing temperatures, than microorganisms profit from a temperature increase; thus, a scenario of modelling fauna at a temperatures increased by 1.5 °C and more so at 3°C will show decreased degradation of organic carbon, leading to higher organic carbon remaining than in the reference scenario with fauna at current temperature.

## 2 Study area and methods

The data that the modelling was based on, were taken from a field study in the Johannisaue, River Fulda floodplain, described e.g. in Husmann and Marxsen (1988) and Marxsen et al. (2021). This study is to this date one of the most comprehensive (chemical and physical conditions, prokaryotes and fauna; microbial growth rate measurements) and longest-running studies (measurements between 1978 and 1984), including dividing prokaryote cells into size fractions allowing detailed biomass distributions. Marxsen et al. (2021) had performed a k-means cluster analysis on the physical and chemical properties, assuming four groups. They characterized the four groups based on the chemical and physical properties of the groundwater sampled in the wells within the groups, and the wells’ position in the floodplain (compare S1: Fig. S1). These groups are used here for different exchange situations at the same climate.

Groundwater temperature was only measured at the time of sampling in the wells in the river Fulda floodplain, i.e. at the maximum monthly. Therefore, we took the daily temperatures from the DWD (German weather service) station Fulda Horas (1526, 255.00 m a.s.l.; geographical latitude 50.5335; longitude 9.6750; on the east shore of the river at the north of the investigated area). From the relationship of the minima and maxima of the *in situ* groundwater temperatures, compared to the minima and maxima of the Fulda Horas air temperature, we derived daily groundwater temperatures separately for the four groups of wells (Supplement S2: Fig. S2). These were the basis for the scenario runs where temperature was one variable influencing maximum microbial growth rate and microbial half saturation concentration (refer to Supplement S2 for further explanations). We used the precipitation data from the same meteorological station 1526: Fulda Horas for estimating recharge (Supplement S3). We then modelled carbon import into groundwater by multiplying the respective daily recharge volume from precipitation with an average carbon concentration. All carbon sources, and all organism compartments, were transformed to mol COD, i.e. chemical oxygen demand, to allow directly comparing compartments. See Supplement S4 for these calculations.

For modelling microbial and faunal growth, we chose hybrid Monod / Verhulst models (Schlogelhofer et al., 2021), combining resource-dependent growth with the limiting factor of a carrying capacity. For microbial growth, the equation reads

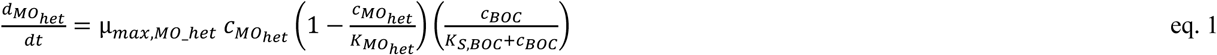

where *c*_BOC_ is the concentration of biodegradable organic carbon (BOC; see below), *c_MO_het_* is the concentration of heterotrophic microorganisms, *µ_max,MO_het_* the maximum growth rate of heterotrophic microorganisms, *K_MO_het_* the carrying capacity, and *K_S,BOC_* the half saturation concentration (Schlogelhofer et al., 2021). We derived growth rate and carrying capacity from the maximum biomass (dry mass) measured in the wells (Supplements S5, S6, S7). Modelled biomasses unlimited by something like a carrying capacity tend to increase exponentially because natural limiting processes such as heterogeneity of the environment are not incorporated in the model. The equation for faunal grazing on microorganisms was built accordingly. For fauna, the model was implemented with *c_MO_het_* instead of *c_BOC_*, *c_fauna_* instead of *c_MO_het_*, *µ_max,fauna_* instead of *µ_max,MO_het_*, *K_fauna_* instead of *K_MO_het_* for the carrying capacity, and *K_S,MO_het_* instead of *K_S,BOC_* for the half saturation concentration.

With the paradigm “Everything is everywhere but the environment selects” (Baas Becking, 1934) still holding (de Wit and Bouvier, 2006), microorganisms can be assumed to change in composition when environmental conditions change. The functionality largely remains the same, due to functional redundancy (Taubert et al., 2019). For microorganisms, there is no mortality in the stricter sense. Instead, maximum growth rate and carrying capacity are parameters usually describing limited growth quite well. Since most bioreactions are most efficient at temperatures between 25 and 35 degrees (Kong et al., 2020), temperature increases in groundwater can be expected to lead to increases in microbial functions up to ca. 35 °C, which will probably never be reached in groundwater unless under geothermal or anthropogenic influences. Therefore, we chose the simplistic approach of modelling temperature increasing microbial growth rate, based on Schmidt et al. (2018). Due to the lack of knowledge on parameters for faunal growth rates, we chose a linearly increasing temperature-induced mortality instead, based on data in Avramov (2013; see below and Supplement S7).

Models were run separately for the four well groups derived in Marxsen et al. (2021). Some characteristics, such as air temperature, precipitation, TOC (total organic carbon) in precipitation, fauna growth parameters maximum growth rate *µ_max_*, half saturation concentration *K_s_*, and carrying capacity for microorganisms and fauna, fauna mortality rate, fauna excretion rate, were the same for all four groups, because group-wise measurements were lacking. The group-wise means of the first date from which measurements were available, for detritus, DOM, heterotrophic microorganisms’ biomass (dry mass), microbial growth rate parameters *µ_max_*, half saturation concentration *K_s_*, and fauna dry mass were taken as the start values for the model.

The values for particulate organics measured in the River Fulda floodplain (Marxsen et al., 2021) were assumed here to be representative for detritus, and it was assumed that continuously, a fraction k1 of 0.0001 per day would become biologically available, and this form was referred to as BOC (biologically oxidizable carbon) here.

Since the remaining BOC concentrations after the modelled period of 3.5 years might just be a hot or cold spot or moment (McClain et al., 2003), we fitted trends over the modelled period to approximate overall development of carbon in the respective compartment. This also allowed to compare fitted trends among scenarios.

With a regional sensitivity analysis (RSA) based on Pianosi et al. (2015, 2016), implemented in the R package “SAFER” v. 1.2.1 (Gollini et al., 2025) parts of which we adapted to the groundwater trophic model, we narrowed down which of the deduced and estimated variables were most influential and warrant special attention in the future. To demonstrate the influence these parameters indeed had on the model, we performed two additional runs for the most influential parameter, with a minimum and maximum value tested for sensitivity, respectively (see model parameters in “parameter_variables.xlsx” in the “gtm” (groundwater trophic model) repository on github: https://github.com/suisch/gtm_River_Fulda_floodplain.).

All simulations, statistical evaluations, and plots were done in R 4.5.0 (R Core Team, 2025). Colours were chosen to be distinguishable by colour blind people using the package “ggokabeito” (Barrett, 2021).

## 3 Results

Precipitation and air temperature showed the expected seasonality, with the years 1978 having been drier, and the year 1981 wetter than the other years (Fig. 1 (a)). Figure 1 (b)-(e) show the measurement points and the simulations results (lines) over the investigated period. Supplement S7: Fig. S3 includes the trends added to Fig. 1 (b)-(e) as an example. The other figures with measured values and trends are not shown because the figures would have been largely redundant with the comparison of trends across scenarios (see below). The trend parameters of all scenarios are given in S10: Table S1.

**Figure 1:**
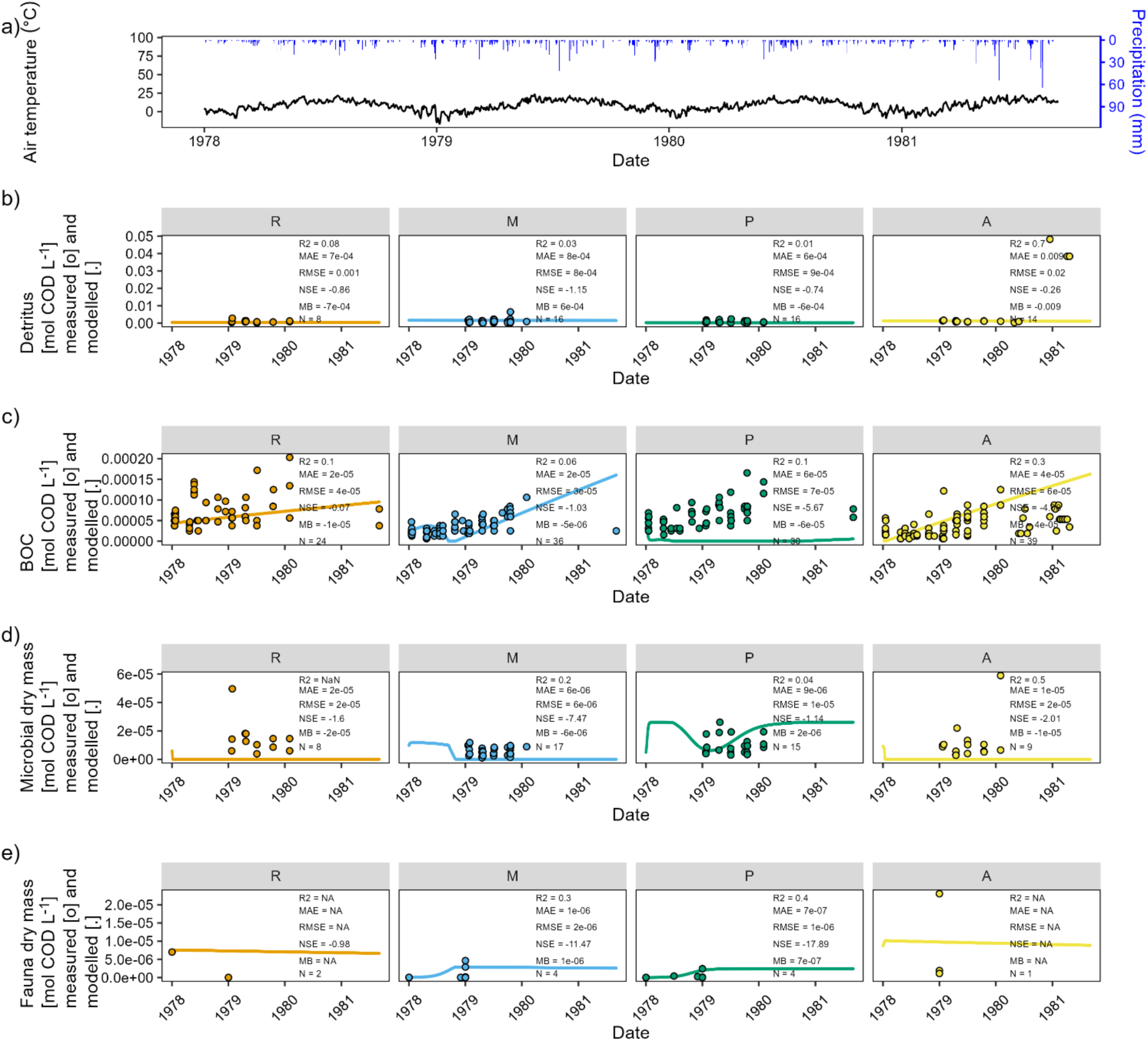
Visualization of the measured and modelled carbon compartments in the river Fulda floodplain for which Marxsen et al. (2021) had divided groundwater wells into four groups: river-near = “R”, mixing zone = “M”, plume = “P”, agricultural area = “A”. (a) Air temperature (black) and precipitation in Fulda Horas meteorological station 1526 (blue) for the four years covering the study period. (b), (c), (d), (e): measured (dots) and modelled (lines) carbon compounds in the four well groups (refer to text and SI Fig. 1). This scenario is referred to as the “Reference” scenario. Please note that the highest measured value in fauna dry mass (1.3*10^−4^ mol COD L^−1^ in P03, group “R”; on 1 January 1979) was omitted from the plot to enable visualization of the modelled time course. In each subplot, 5 error measures and the number of values between measured data averaged per group and date, and the model data are given.

The simulations (lines) in Fig. 1 (b), (c), (d), and (e) reflect the measured values (points) well in general. We therefore used this model and changed only the parameters for the presence of fauna and for temperature in the scenarios of “no fauna” and / or “+ 1.5 °C”, and / or “+ 3 °C”, respectively (described below). The respective overviews of the same format as Fig. 1 are included in the Supplementary Information S8: Fig.s S3 to S8.

The modelled detritus did not change much during the modelled time span (Fig. 1 (b)). Event-based recharge in response to precipitation seemed to contribute little (Fig. 1 (b)). Modelled BOC first decreased then increased in the agriculturally-influenced “A” and the plume “P” zones (Fig. 1 (c)). In the mixing “M” zone, after an initial increase, there was a decrease in BOC until autumn 1978 and then a steady increase. In the mixing “M” and the agriculturally-influenced “A” zones, BOC reached higher values at the end of the modelled time span than the measured values (Fig. 1 (c)). The increasing trend was largely consistent with an apparent increasing trend in the *in situ* values over the study period (Fig. 1 (c)).

The microbial dry mass initially decreased in the “R” and “A” zones and stayed at low levels (Fig. 1 (d)). In the “M” and “P” zones, dry mass initially increased to the carrying capacity, which was set to be the maximum microbial dry mass measured per group. The dry mass more or less stayed on the carrying capacity or slightly below, for the better part of 1978, and then decreased. This decrease was only temporary in the “P” zone (Fig. 1 (d)).

Modelled fauna dry mass (after an initial slight increase in zone “A”) slowly and steadily decreased in the river-near “R” and the agriculturally-influenced “A” zones (Fig. 1 (e)). In the “M” and “P” zones, fauna dry mass increased until late autumn 1978, and then stayed on a level just below the carrying capacity (the highest measured fauna dry mass per group).

The modelled BOC, microbial dry mass and fauna dry mass showed only slightly different patterns in the “+1.5 °C” scenario (Fig. S5), but very different patterns in the “+3 °C” scenario (Fig. S7). The three “no fauna” scenarios (Fig.s S4, S6, S8) showed seasonal patterns for faunal dry mass. At +1.5 °C, fauna decreased seasonally, presumably because in summers, the temperature became lethal. However, this did not seem to influence net microbial growth and net BOC degradation. At +3 °C, however, fauna seemed to break down to low levels in summer 1978, from which they hardly recovered in the following springs and stayed low afterwards. Due to the lacking faunal grazing, microbial dry mass reached highest values, even to the parameterized carrying capacity. In turn, the resulting BOC values after the 3.5 years were the lowest of all scenarios, particularly in the “R” and “P” zones.

An overview over the start and end values of the modelled run, and of the linear fit, along with the fit parameters, is given in S10, Table S1. The comparison of all trends is given in Fig. 2 for BOC, for microbial dry mass in S9, Fig. S9, and for fauna dry mass in S9, Fig. S10. In all zones, the three “no fauna” scenarios (“No fauna”, “No fauna +1.5°C”, “No fauna + 3°C”) were indistinguishable from each other (yellow, bright orange and dark orange lines fall together). For the “M” and “P” zones, BOC trends rose to the highest values per zone, thus, highest BOC values remained in the system in the “no fauna” scenarios in these two groups (Fig. 2). Except for the “+3°C” scenario in the “R” group and all “fauna” scenarios in the “P” group, BOC trends rose in all scenarios. In all four groups, the BOC trends for the “reference” and “+ 1.5°C” scenarios were indistinguishable from each other. In the “P” zone, the “+ 3°C” scenario also fell visibly into one line with the two other “fauna” scenarios. The main difference was thus not among scenarios of varying temperature, but among “fauna” and “no fauna” scenarios (Fig. 2).

**Figure 2:**
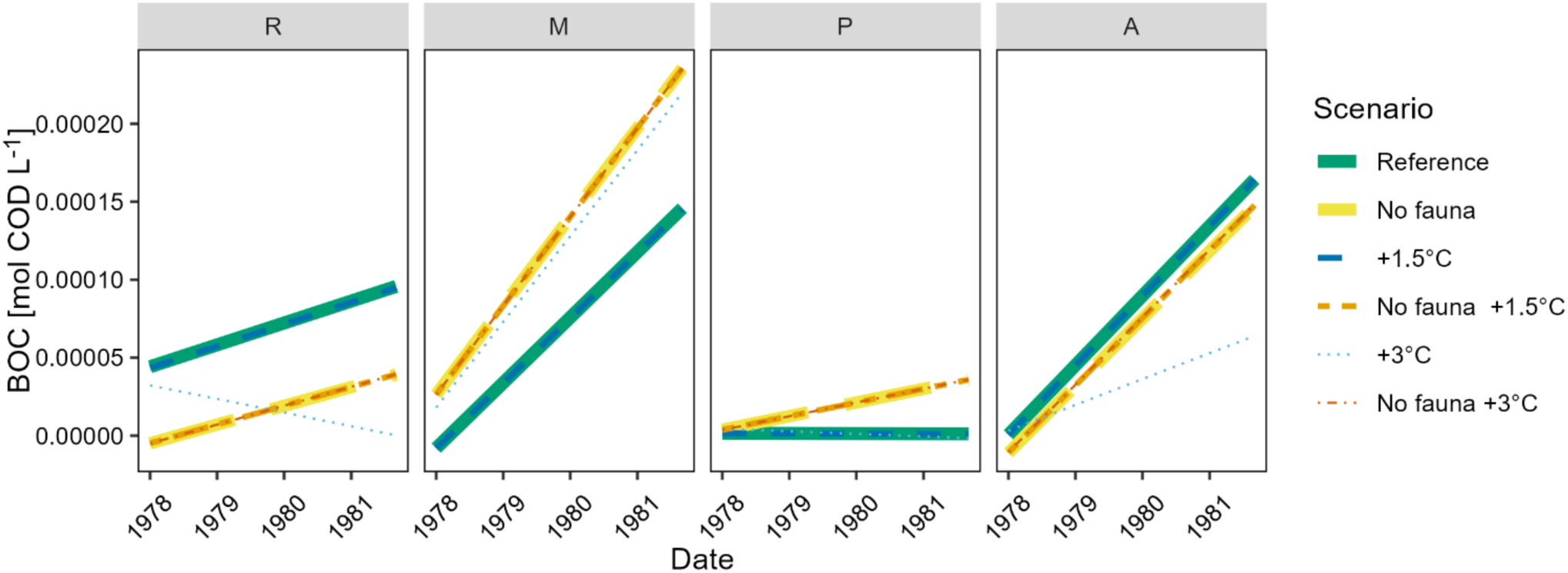
Trends through the modelled BOC in the river Fulda floodplain for the six scenarios which are given in the legend: “Reference”: input variables according to Marxsen et al. (2021); “No fauna”: setting all fauna variables to 0; “+1.5°C”: raising all groundwater temperatures by 1.5°C; “No fauna +1.5°C”: setting all fauna variables to 0 and raising all groundwater temperatures by 1.5°C; “+3°C: raising all groundwater temperatures by 3°C; “No fauna +3°C”: setting all fauna variables to 0 and raising all groundwater temperatures by 3°C. The four sub plots depict the four aquifer groups: river-near = “R”, mixing zone = “M”, plume = “P”, agricultural area = “A”; see text and SI 1.

The ecosystem service of carbon degradation was expressed as the concentration of mol COD of biologically oxidizable carbon BOC remaining in the system compared to the start value of the simulation, with column “Modelled concentration at end minus modelled concentration at time 0” being the difference between the columns “Modelled concentration at the end” and “Modelled concentration at time 0” (S10, Table S1). We divided this difference by the value in column “Modelled concentration at time 0” to yield the value in the column “% change at end compared to modelled concentration at time 0”. BOC decreased by almost 100% over the modelled period to very low values in the fauna scenarios in the plume “P” zone (Supplement S10, Table S1), while it increased by over 700% during the same period in the “no fauna” scenario in the mixing “M” zone. This shows that it depends on the start condition and measured microbial rates per zone what the outcome of the simulation will be – there is no general behaviour. We then related the “Modelled concentration at the end” of any scenario to the reference case to yield “factor by which modelled end concentration changed from reference modelled end concentration”. A value below 1 indicates a decrease in the respective concentration compared to the reference case. For BOC, this was observed in the “R” and “A” zones without fauna, and in the “R” and “P” zones at elevated temperatures, and at +3°C additionally in the “A” zone (S10, Table S1). In the “+1.5°C” and “+3°C” scenarios, the remaining BOC increased only slightly above the reference level: with a factor of 1.37 in the “M” zone at +3°C. In the “no fauna” scenarios, the remaining BOC increased by a factor of up to 6.6 in the plume “P” zone, compared to the reference scenario, regardless of the respective temperature increase (S10, Table S1).

In partial agreement with the second hypothesis, an increase in temperature by 3°C led to lower remaining BOC in most scenarios. However, an increase of 1.5°C did now show a visible effect, except for the fauna dry mass. Microbial dry mass fitted trends increased beyond the reference case in the “no fauna” scenarios, except for the “P” zone. Fauna dry mass showed lower fitted slopes for the “+3 °C” scenarios in the “R” and “M” zones. The trends of faunal dry mass at “+3°C” was different in every zone and (almost) fell together with the “no fauna” scenarios in the “P” zone. Clearly, a groundwater temperature increase by 3°C had impacts on fauna which in the “R” and “A” zones led to increased microbial dry masses, which, in turn decreased remaining BOC trends in the “R” and “A” zones. The increase of temperature, however, cannot be assumed to act on all aquifers in the same way.

In order to deduce how sensitive the model was to changes in specific parameters, we conducted a regional sensitivity analysis separately for the four zones. In all zones, k1, i.e. the proportion with which detritus split into biologically oxidizable carbon, was highly sensitive (S 11; Fig.s S11 – S14).

Since microbial yield on carbon, proved so sensitive, we ran the reference scenario with the minimum (0.001; Fig. S15) and maximum (0.8; Fig. S16) microbial yield tested in the sensitivity analysis. Low yield and high grazing pressure from fauna led to microbial dry mass declining to lowest values (Fig. S15). BOC remaining in the system was high. High yield, in contrast, increased microbial growth in the “M” and “P” groups to carrying capacity, and an initial peak in the “A” group (Fig. S16). Fauna dry mass was higher in the “M”, “P”, and “A” zones than in the low microbial yield scenario. Resulting BOC was higher in the low yield scenario than in the high yield scenario.

## 4 Discussion

This is to our knowledge the first attempt at developing a process-based model of the groundwater food web, and it is the first endeavour at comparing scenarios of excluding fauna from the system. There have been laboratory studies exposing fauna to increasing temperature (e.g. Brielmann et al., 2011; Di Lorenzo and Reboleira, 2022; Spengler, 2017), but to our knowledge this has never been translated to a trophic dynamic model, even if restricting to the major part of the food web, as done here. One reason for such attempts not having been done before, lies in the difficulty of parameterizing such models. Our knowledge of the groundwater ecosystems’ processes and dependencies is still patchy. Thus, the choice and development of appropriate parameters is crucial. If the reference case is not appropriately parameterized, using it as a base for further scenarios is not reliable.

### 4.1 Modelling the reference case with fauna at the temperature of the observation period

Largely, the model set up, including the parameters, proved to be appropriate for representing the measured data. Modelled detritus in the reference case with fauna at the temperatures of the observation period stayed in the same order as most of the measured values and was a constant source of carbon in abundance. We assumed here that detritus would largely stay in the catchment. We further assumed that BOC would be the more influential factor for food webs and drinking water production, as e.g. in Vogt et al. (2023), and thus, we focussed on BOC and its pathway in the following.

Modelled BOC concentrations, designed to represent the bioavailable carbon fraction, mostly increased, and even beyond the measured values, over the investigated period of 3.5 years in the mixing “M” and the agricultural “A” zones. The model thus did not indicate a quasi steady state. This may be an artefact of the combination of model parameters and needs to be studied further. However, this increase was in the order of 0.00015 mol COD L^−1^ over the investigated period of 3.5 years, while the highest values of detritus measured in the field were around 0.05 mol COD L^−1^. Thus, this BOC increase was negligible for the catchment-scale budget. It just means that, seen over the whole 3.5 year modelling period, less of the biologically available carbon pool was taken up by microorganisms than was refilled from precipitation, physical breakdown, and import from the soil. Assuming that the ecosystem in fact should have been in a quasi steady state, the trends would have been expected to be even closer to 0 in the reference model case, but for a simplistic food web model, we believe that the current approach is sufficiently close to the measured values. It further ensured that carbon was not limiting at the bottom of the food web over the whole period. This enabled comparison of the degradation of this resource being in over abundance. The less exchange with the surface, i.e. the deeper and the more secluded aquifers are, the more dependent on carbon import they will be (Schmidt et al., 2017; Schmidt and Hahn, 2012). Such scenarios will be part of the next questions to be addressed with the model.

In the scenarios, temporarily, modelled BOC was reduced to near-zero during the growth phase of the microorganisms (see below) in the mixing “M” and the plume “P” zones. This indicates feedback among the carbon fractions’ concentrations – their scales seemed to be appropriate to reflect dynamic ecosystem behaviour. It also means that a longer-term simulation might have yielded different end concentrations and trends. This may be the subject of further model applications.

Modelled microbial dry mass showed (at least temporary) reductions to near-zero concentrations. The early quite sudden dips in the “A” and “R” zones might reflect moments of feeding by higher-than-average fauna biomass (see below). The highest measured value for microbial dry mass had been observed in the agricultural “A” zone (compare Marxsen et al., 2021) and concurrently, carrying capacity was set to be at this high level, but was never nearly reached in the model with the current parameter set up. One reason for this may be that the modelled fauna dry mass stayed highest in the “A” zone compared to other zones and thus, fauna exerted grazing pressure, in this case reducing the microorganisms’ capacity of degrading BOC, such as observed in Kota et al. (1999). This can be seen in the steadily rising modelled BOC. It is likely that the groundwater microbial community under agricultural land use exhibits growth and degradation characteristics much different from other zones (e.g. Dibbern et al., 2014), because subsurface carbon characteristics differ under the different land uses (Lehmann et al., 2021). It might be necessary to adapt the parameters and maybe even the model set-up for agricultural zones in future work. Another source of variation may be the distribution of organic carbon within the aquifer – in a porous environment, it might be physically separate from the microbial cells (Schmidt et al. 2017). To take such micro scale resource heterogeneity into account, requires a model which parameterizes at least two spatial dimensions (Schmidt et al. 2018), instead of the spatial 0D approach taken here.

Modelled fauna dry mass showed a steady increase within the first year in the mixing “M” and plume “P” zones, concurrent with a decrease in the microbial dry mass in the same zones, particularly towards end of winter in early 1979. With the end values being lower than the start values in the “R” and “A” zones, the fitted trends across the models were negative. The fauna fitted trends for the “+3°C” scenarios decreasing to below 0 are an artefact. In the cases where the dry mass stayed at elevated levels over at least a certain period at the start, the drop down was steeper than in scenarios where fauna decreased at the beginning. The reason for the fauna dry masses not increasing later when the microbial dry mass was at carrying capacity, i.e. when food was presumably not limiting, e.g. in the “+3°C” scenario, requires further investigation, but is most likely due to increased mortality of fauna at generally higher temperatures. Experiments on feeding and growth rates, building up on the published experiments on reproductive rates in Rütz et al. (2022) from central Germany, are still scarce and hardly available for middle-European temperate weather groundwater environments.

We had checked data from three studies for their potential use as parameters in the current model. These experiments had been conducted at 14 and 17°C (Di Lorenzo et al., 2025), at 15°C (Navel et al., 2011), and at 10°C (Mermillod-Blondin et al., 2025). Thus, only the study by Mermillod-Blondin et al. (2025) was under conditions somewhat comparable with the River Fulda floodplain. The study by Navel et al. (2011) was even conducted with a 12 h light 12 h dark cycle because of the studied interaction with a surface amphipod. Both elevated temperature (see above) and light (Simčič and and Brancelj, 2007) may be stressful for stygobites though, and we did not deem the parameters appropriate for our model. This reduced the available parameters. Considering these caveats, for the sake of a first simplistic food web model, we considered the model set-up sufficient for the purpose.

### 4.2 Modelling the lack of fauna

In partial agreement with the first hypothesis, the “no fauna” scenarios in the “M” and “P” zones resulted in the highest remaining BOC, both in models, and in the fitted trend lines. In parallel, microbial dry mass was mostly highest in the “no fauna” scenarios. Less microbial degradation despite higher measurable dry masses points to a lack in rejuvenating grazing (Mattison et al., 2005). The bulk growth of microorganism was probably higher in the “fauna” scenarios, leading to higher degradation, but faunal grazing reduced the measurable microbial net dry mass (see also above). Elucidating the actual microbial growth and uptake of organics per cell requires tracer studies (e.g. for marine systems: Zubkov & Tarran, 2008). It is interesting that this pattern emerges clearly from the rather simple food web modelling with the hybrid formula of combining the Monod model with the Verhulst model (Schlogelhofer et al., 2021).

The opposite effect of microbial uptake being reduced due to faunal grazing, described by Kota et al. (1999), was observed in the “R” and “A” zones. We lack understanding of the circumstances under which one pattern dominates. Also, the cited publications refer to grazing by protozoa, while we lacked quantitative field data for protozoa from the River Fulda floodplain and assumed that the grazing can be subsumed under a multicellular “fauna” dry mass term, based on the fauna appraisal in the River Fulda floodplain. To date, there is no field or laboratory study quantifying multicellular fauna, protozoa, and microorganisms in the same groundwater samples. Qualitative food webs were constructed based on molecular data in the Hainich aquifer (Herrmann et al., 2020).

Depending on the zone, remaining BOC concentrations decreased to the point of vanishing (plume “P” zone in the “+ 3°C” scenario; discussed below), but increased in all other cases. This increase was up to more than 6.6-fold over the 3.5 year observation period. Without groundwater fauna, drinking water production will have to compensate by intensified treatment of organics which raises costs and insecurity.

### 4.3 Modelling an increase of groundwater temperature

It was surprising that an increase by 1.5°C, while resulting in different fauna dry mass (except for the “P” zone), did not translate in different microbial or BOC fitted trends in the “fauna” scenarios. This may show that the lower grazing pressure from fauna was apparently compensated by faster microbial growth resulting in the net biodegradation of BOC being no different from the reference scenario. It might also have been a combination of the effect described by Kota et al. (1999) and thermally increased biodegradation rates. Fauna being clearly compromised by +3°C translated into higher microbial dry mass. This higher microbial dry mass, however, did not alter BOC fitted trends in the “P” zone. Here, the diminished “rejuvenating” grazing might have compromised degradation, along with Mattison et al. (1999). The strongest increasing by 3°C effect on faunal dry mass was seen in the “R” and “A” zones. In both zones, microbial dry mass fitted trends steeply increased over the course of the 3.5 modelled years, almost in parallel. However, the BOC fitted trends in these two zones were not parallel: it decreased in the “R” zone and increased in the “A” zone. In the “R” zone, thus, more remaining microbial dry mass degraded more, while in the “A” zone, it might have been a lack in rejuvenating grazing (Kota et al., 1999) again, which compromised degradation.

### 4.4 Modelling the lack of fauna at an increase of groundwater temperature

In contrast to the third hypothesis, the combination of increased temperatures and lack of fauna led to reduced degradation and higher remaining BOC compared to the reference scenario, only for the “M” zone at +3°C. In that scenario, the difference to the BOC increase in the “no fauna at recent temperature” was negligible and thus, the reduced degradation was almost only due to lack of faunal grazing. In the “P” zone, the only factor shaping the BOC fitted trend was presence versus absence of fauna – temperature did not influence at all. The lack in temperature effects on fauna might partly be due to the overall low dry masses of fauna in the “P” zone.

While the faunal and microbial dry mass fitted trends were similar in the “R” and “A” zones, the BOC fitted trends diverged, particularly for the “+3°C with fauna” scenarios. The reason may lie in different indirect effects which are hard to capture with the present set-up. More combinations of start conditions and parameters are needed to single out direct and indirect effects (Fišer et al., 2025).

BOC trends were lowest for the “+ 3°C” scenarios in the “R” zone, and thus, the lowest BOC remained at the highest temperature with fauna, although fauna was clearly compromised. One explanation is that the “intermediate” disturbance caused by the temperature did, in this case, not lead to the highest diversity, as proposed in the original “intermediate disturbance hypothesis” (Connell, 1978), but to the highest ecosystem services. However, it needs to be tested how sustainable this pattern would be in the longer run under increased temperatures.

In general, the effect of losing fauna was mostly stronger than that of increased temperatures. The risk of losing fauna altogether increases with increasing temperatures, and thus, the risk of running into “no fauna” scenarios with partly clearly decreased degradation, increases.

### 4.5 Reduced ecosystem services

Here, we focused on degradation of BOC as a main ecosystem service relevant to drinking water production for humankind. In many scenarios, including the “no fauna” scenarios of the “M” and “P” zones, BOC concentration increased, i.e. the remaining carbon increased, which is an indicator of a reduced ecosystem service. Thus, the ecosystem service was reduced mainly under the lack of fauna. Losing groundwater fauna would mean losing a substantial part of the microbially-driven ecosystem service of carbon degradation.

This ecosystem service of degrading organic carbon comes at costs. E.g. the oxidation of carbon to CO_2_ requires oxygen. There are a few laboratory studies quantifying the uptake of oxygen by groundwater fauna, e.g. Di Lorenzo et al. (2024), Hervant (1997), Hervant et al. (1998, 1999), Malard and Hervant (1999), and Mermillod-Blondin et al. (2008). Future studies should make dissolved oxygen a reaction partner in the organisms’ uptake models, in order to evaluate where the provision of ecosystem services may be hampered by the aquifer’s oxidation state. The wells in the Fulda aquifer which we used to parameterize the current models were largely fairly well oxygenated and dissolved oxygen was shown to be not limiting in Marxsen et al. (2021), but in other aquifer situations, dissolved oxygen played a major role in shaping communities (Strayer et al., 1997) and thus needs to be included in future models which should be applicable for other areas as well.

### 4.6 Sensitivity analysis

The two parameters showing higher sensitivity in one group each than other parameters were the microbial growth parameter yield and the k1 (proportion with which detritus split into biologically oxidizable carbon). It is thus paramount to derive these two parameters carefully, to exclude artefacts, since we are showing here that changing microbial yield may change the outcome of the model considerably, particularly for BOC. It needs to be noted though that particularly facultative oligotrophs are known to exhibit different growth parameters when facing different substrate concentrations (Ishida et al., 1982). This will be hard to parameterize representatively and questions how important exact microbial growth parameters can be in reality.

### 4.7 Full budget

Full accounts of the productivity of groundwater systems with or without fauna, and at different temperatures, require incorporating CO_2_ into the budget. While dissolved bicarbonate had been measured in the River Fulda aquifer study, some of the CO_2_ (and CH_4_ if produced; Herrmann and Taubert, 2022) will gas out. Such carbon loss from the system needs to be measured with the appropriate methods. Isotopes of the carbon atom may assist in such calculations (Schwab et al., 2019). A study on all carbon fractions in a groundwater ecosystem is still lacking, – the AQUADIVA project (Küsel et al., 2016) offers probably the most complete data set on all compartments. But important grazer groups such as the rotifera (Ejsmont-Karabin and Karpowicz, 2025) and protozoa (Sinclair et al., 1993) require different sampling methods than the larger fauna and are thus often not included in quantitative groundwater grazer accounts. For a full budget of the standing stocks and processes in the groundwater ecosystem, these compartments need to be included in future field studies and in the models based on their results.

### 4.8 Extrapolating to the global scale

The total groundwater volume in the upper two km of the continental crust was modelled, based on tritium, to be approximately 22.6 Mio. km^3^ (Gleeson et al., 2016). Of these, Gleeson et al. (2016) estimated that between 0.1 and 5 Mio. km^3^ are less than 50 years old. Abbott et al. (2019) drew the conclusion that between 300 and 1200 km^3^, with an average of 630 000 km^3^, are renewable groundwater (“young” and mostly fresh). This “younger” groundwater is often the resource used for drinking water production.

We found increases in the BOC concentrations from the reference scenario to increased temperature and no fauna scenarios, with the increase being over 6.6-fold in the case of the “no fauna” scenarios in the mixing zone “M” group, which was a difference of 7.3*10^−5^ mol L^−1^ over the observation period of 3.5 years, i.e. 2.1*10^−5^ mol L^−1^ year^−1^. Assuming that this difference would be representative for global conditions, multiplying with the average of 630,000 km^3^ for renewable groundwater (Abbott et al., 2019) and with a molar mass of 227 g mol^−1^ for humic acid (see Supplement S4), this translates to globally, up to 2,982,780,000 tons year^−1^ not degraded by the groundwater ecosystem, when fauna is lacking. This is an additional load of COD that drinking water production needs to reduce globally in the worst case when and if groundwater fauna has become extinct.

Extrapolating like this assumes that the situation in the Fulda floodplain is representative for the whole world, which is unlikely to hold true, for various reasons. One step in making such scenario accessible for the global situation is to use spatially resolved temperature scenarios such as by Benz et al. (2017, 2024) to extrapolate groundwater ecosystem services to the national, continental, and global scale. Another important step is to make not only the microbial degradation rate dependent on temperature, but also the fauna grazing rate, not only the fauna mortality.

### 4.9 Necessary steps for further development

In this modelling study we considered different kinds of energies as equal, due to a lack in better knowledge and data. This approach may lead to errors (Odum and Odum, 2000). Odum and Odum (2000) suggested to express everything in an ecosystem in the solar energy used to make each item. However, in an ecosystem where the sun does not shine, and where instead not only the import of plant material, but also chemolithoautotrophy without solar energy provide resources at the basis of the food web (Herrmann et al., 2015; Kellermann et al., 2012), compromises must be found.

Many temperature-related processes are not considered here, namely environmental changes, such as lower solubility of dissolved oxygen, higher dissolution of ions (Retter et al., 2020), all of which may be detrimental to organisms. Thus, the net optimum for groundwater microbial activity is probably lower than the physiological activity optimum itself. However, we lack parameters for groundwater ecosystems to include such dependencies into models.

This study was based on a plethora of assumptions most of which might trigger discussions. Also, here, only average compartment carbon concentrations were modelled. It is relevant to focus on the whole range of likely conditions, i.e. including minima and maxima. However, because of error propagation, this is not a straightforward process and requires diligent development. The present study aims to serve as a starting point for future developments.

Climate change does not happen everywhere in the same way – it not only translates into increased temperatures, but also into changed recharge patterns which were not considered here. One next step is to use spatially resolved temperature scenarios and chemical reaction solvers such as PHREEQ-CE to extrapolate groundwater ecosystem services under climate scenarios to the national, continental, and global scale.

## 5 Conclusion

This first study on quantifying the change in ecosystem services at scenarios of losing groundwater fauna, and of increased temperature, showed that the partial loss of ecosystem services (BOC degradation) due to lacking fauna increased remaining BOC up to 660-fold, based on the 3.5-year observation period. While there were differences among aquifer zones that were characterized by differing patterns in nutrient and carbon background, some general patterns emerged. This means that biomanipulation by protecting and promoting groundwater fauna for ameliorating water quality, might be a promising concept. The fauna’s role might still be underestimated. This has implications for drinking water production which will locally have to compensate the lack of fauna’s ecosystem services by increased treatment which raises costs and insecurity.

## Appendices

The supplement related to this article is available in the file SI.pdf.

## Code availability

Code for a former version of the model and for producing the results graphs is available on zenodo at https://doi.org/10.5281/zenodo.17153927 (Schmidt, 2025). This is equivalent to v.1.0.0 of the github repository https://github.com/suisch/gtm_River_Fulda_floodplain. For this contribution, this repository has been updated to v.2.0.0.

## Data availability

Parts of the data from the River Fulda floodplain study had been published in Marxsen et al. (2021). Data are available on zenodo at https://doi.org/10.5281/zenodo.17153927 (Schmidt, 2025) and on the github repository https://github.com/suisch/gtm_River_Fulda_floodplain. For this contribution, this repository has been updated to v.2.0.0.

## Supplement link

The supplement file is available under the link

## Author contributions

SIS conceptualized the study and prepared the methodology of the model. SIS, JM, and NR curated the data. SIS, JM, and NR wrote the original draft. SIS, JM, and NR validated the results. SIS wrote, reviewed and edited the final version.

## Disclaimer

Copernicus Publications remains neutral with regard to jurisdictional claims made in the text, published maps, institutional affiliations, or any other geographical representation in this paper. While Copernicus Publications makes every effort to include appropriate place names, the final responsibility lies with the authors. Views expressed in the text are those of the authors and do not necessarily reflect the views of the publisher.

## Acknowledgements

SIS thanks Carola Winkelmann, Thomas Petzoldt, Martin Thullner, Jan Ulrich Kreft, Muhammed Shikhani, Martin Schultze, and Susanne Worischka for discussions on modelling.

