## Supplementary material for "Quantifying the impact of groundwater fauna and temperature on the ecosystem service of microbial carbon degradation": Supplmental figures and tables

### S1: Map of the study region and situation of the wells and weather station

The map with the locations of the wells and their respective position in the aquifer zones derived in Marxsen et al. (2021) are shown in Fig. S1.

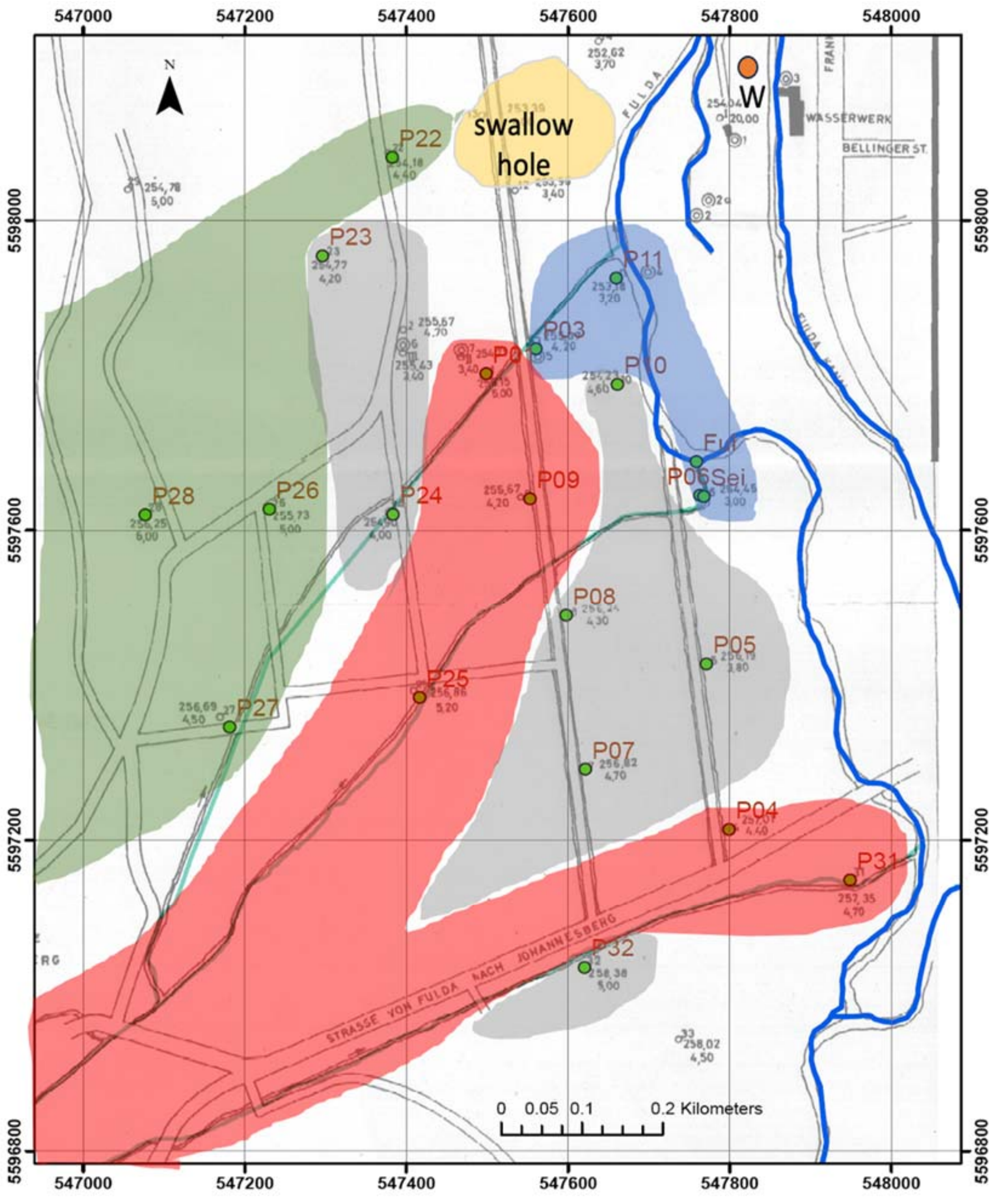


*Figure S1: Map of the study area near Fulda, with the four groups of wells derived in Marxsen et al. (2021) colour-coded. Blue: river-near, abbreviated as "R", grey: mixing zone ("M"), red: plume ("P"), and green: agricultural area ("A"). W: weather station Fulda Horas, DWD 1526. Changed after Marxsen et al. (2021). Coordinate system: ETRS 1989, UTM Zone 32N.*

### S2: Derivation of the daily groundwater temperatures in the four well groups

The daily temperature series (daily measurement of the average air temperature at 2 m a.sl. in °C) from DWD (German Weather Service Wetterdienst (DWD), accessed 10th of April 2025) were scaled to groundwater, according to the minima and maxima of the *in situ* measurements in groundwater and the minima and maxima of the air temperature values during the same period (Fig. S2).

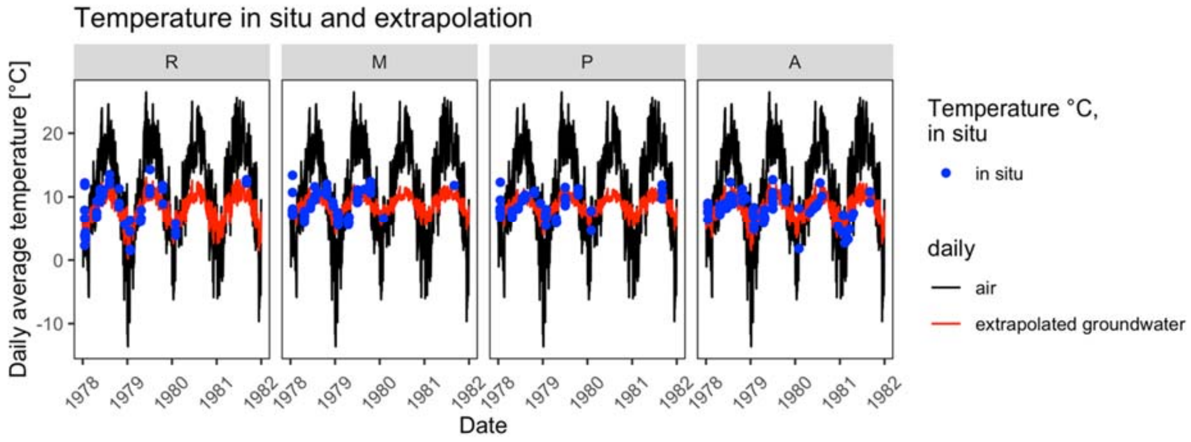


*Figure S2:* In situ *groundwater (blue dots), daily measured air temperature in Fulda-Horas (black line; position see map in Fig. S1), and extrapolated daily groundwater temperature (red line), in the four well groups (see Fig. S1).*

### S3: Estimating carbon recharge from precipitation

There are surprisingly few data on DOC or TOC concentrations in precipitation, and even fewer site-scale budgets are available. One such data set is from the Plešné and Čertovo catchments, two areas within 50 km from each other, where the average precipitation in mm per year was 1419.5 mm $year^{-1}$, i.e. L $m^{-2}$ $year^{-1}$ (Kopáček et al., 2009). The TOC in mol $m^{-2}$ $year^{-1}$ was on average 0.2035 (Kopáček et al., 2009). Dividing this TOC load by yearly precipitation yields an average concentration of 0.14 mol $m^{-3}$, i.e. 0.00014 mol $L^{-1}$. For lack of more resolved data, we assumed that such value is realistic for the Fulda floodplain precipitation as well. Assuming that this TOC can be summarized as humic acid (Saccò et al., 2020), the mol COD equivalent was calculated to be 7.5 times the concentration of mol TOC $L^{-1}$, i.e. 1.075 mol COD $m^{-3}$, i.e. 0.0011 mol COD $L^{-1}$. It was further assumed that 1) we can approximate recharge by multiplying precipitation with a fraction factor of precipitation recharged to groundwater. This depends of course on vegetation, the soil and aquifer matrix and many other factors (Reinecke et al., 2021). Based on infiltration having been found to be 6–13% of the annual precipitation in loess aquifers (Liu et al., 2024) and ca. 8% across Europe (Seidenfaden et al., 2023) we applied a factor of 0.08. We further assumed that 2) recharging precipitation mobilizes organic matter from the soil, i.e. detritus from vegetation which had fixed $CO_2$ from the atmosphere, and that this concentration is considerably higher than that recharged, i.e. the carbon import from precipitation was multiplied by 10 to yield detritus present in shallow groundwater. More precise values were not available for the study area.

This concentration was multiplied with the actual volume of daily recharge multiplied with an assumed depth of aquifer of 10 m, assuming simplistically an immediate mixing throughout the aquifer depth which is of course only a rough approximation. For each date, the result was thus mol COD $m^{-3}$ in the aquifer. In the simulation run, the respective daily mol $m^{-3}$ was added to the mol $m^{-3}$ being left from the previous time step and translated to COD $L^{-1}$ because all other concentrations were modelled as mol COD $L^{-1}$.

### S4: Derivation of the input parameters and variables as mol COD $L^{-1}$

Total prokaryote cell numbers and fauna abundances were taken from Marxsen et al. (2021) and were averaged for each of the derived groups of wells. Total prokaryote cell numbers were expressed as $10^{-6}$ $mL^{-1}$. Dry mass was calculated from these cell numbers according to Marxsen et al. (2021). Across all samples, on average, the prokaryote dry mass was calculated to have been around 200 µg $L^{-1}$.

To calculate a budget from the different types of carbon compounds (prokaryote dry mass, fauna dry mass, DOM (dissolved organic matter), TOM (total organic matter), were translated to mol COD $L^{-1}$.

The microorganism growth parameters were based on the reasoning for the degradation of acetate, a universal compound (see also below). Similar growth parameters on groundwater humic substance could not be found and we assumed that the acetate parameters would be similar to those for humic acids, until we have better values. The COD equivalents of the oxidation of acetate can be seen in the reaction $C_2H_4O_2 + 2O_2$ -> $2CO_2$+$2H_2O$ where 2 mol of $O_2$ are needed to oxidize one mol of acetate, thus COD (acetate) = 2 mol $L^{-1}$.

COD of the more complex organic fractions with less obvious stoichiometry was calculated using the formula

$$\mathrm{C}_n\mathrm{H}_a\mathrm{O}_b\mathrm{N}_c + \left(n + \frac{a}{4} - \frac{b}{2} - \frac{3}{4}c\right)\mathrm{O}_2 \rightarrow n\mathrm{CO}_2 + \left(\frac{a}{2} - \frac{3}{2}c\right)\mathrm{H}_2\mathrm{O} + c\mathrm{NH}_3$$

(https://en.wikipedia.org/wiki/Chemical_oxygen_demand; accessed 1 February 2025).

It was assumed that biomass can be summarized with an average composition $C_1$ $H_{1.8}$ $O_{0.5}$ $N_{0.2}$ (Heijnen, 2006), i.e. a molar mass of 24.6 g $mol^{-1}$. Thus, the stoichiometry for the complete oxidation of biomass was derived with the equation above as $C_1$ $H_{1.8}$ $O_{0.5}$ $N_{0.2}$ + 1.05 $O_2 \rightarrow CO_2 + 0.6\ H_2O + 0.2\ NH_3$. This means that 1.05 mol of $O_2$ are needed for the oxidation of 1 mol biomass, thus 1.05 mol COD $mol^{-1}$ biomass.

TOC in precipitation and organic substance (see below) in the Marxsen et al. (2021) study were assumed to consist largely of humic acids (Vogt et al., 2023). According to ChemSpider, humic acid may have the average molecular formula $C_9H_9NO_6$, and thus, an average mass of 227.172 g $mol^{-1}$ (ID: 32820151; https://www.chemspider.com/Chemical-Structure.32820151.html; accessed 1 February 2025). Applying the above formula to humic acid with the parameters n = 9, a = 9, b = 6, c = 1 yields that oxidizing one mol of humic acid requires 7.5 mol of $O_2$.

The OS (organic substance) values in Marxsen et al. (2021) were assumed to be representative of TOM (total organic matter; or detritus) being recharged into groundwater through the soil passage. For transforming to mol COD, it was assumed that this TOM can be summarized as humic acid, in the same way as TOC from precipitation. Using the same formula as for humic acid in precipitation, mol COD $L^{-1}$ of humic acid was calculated to be 7.5 times the concentration in mol $L^{-1}$.

The COD concentration (g $L^{-1}$) from Marxsen et al. (2021) was taken as the dissolved fraction of recharge, in parallel to the largely particulate OS. Mol COD $L^{-1}$ was derived by dividing by the mol mass of $O_2$: 32 g $L^{-1}$. These concentrations were assumed to be those that microorganisms grow on.

Averages of all these molar concentrations of COD of BOC were calculated for the four groups identified in Marxsen et al. (2021) to calculate four different types of groundwater aquifer situations. These average concentrations were taken as the starting values of the simulations.

### S5: Derivation of the temperature influences on the maximum microbial growth rate and microbial half saturation concentration

The groundwater temperature derived in SI 2 was one parameter influencing maximum microbial growth rate and microbial half saturation concentration. This derivation was based on Schmidt et al. (2018) who had done a linear regression of the optimal specific growth rates of nine species of bacteria versus their optimal growth temperature, based on data in Mohr & Krawiec (1980), covering  a temperature range of 14-44°C. The equation of this linear regression of normalized specific growth rates versus temperature was of the form 0.0304·°C – 0.3255, with an $R^2$ of 0.89. Thus, the drop in growth rate per drop in °C temperature was by the factors 0.9696 for one °C, 0.848 for 5 °C, 0.696 for 10 °C and so on. This equation was used to calculate the growth rate of the heterotrophic microorganism in the present work based on the respective daily average groundwater temperature per area (compare S2, in particular Fig. S2). As explained in Schmidt et al. (2018), the half-saturation constant ($K_{S,MO\_het}$) was scaled by the same temperature factor as the specific growth rate ($\mu_{max,MO\_het}$). We assumed that growth yields were not dependent on temperature. These adjustments were applied to derive the temperature-adjusted kinetic parameters in S4.

### S6: Deduction of further variables and parameters

The time step for the simulations were set to daily. Thus, where possible, e.g. for temperature and precipitation, daily input variables were used and prepared.

Except for temperature, fauna abundance, the chemical properties and microorganism numbers measured in the field, the six scenarios (1: reference conditions, 2: no fauna, 3: temperature elevated by 1.5 °C (refer to SI 2 for the derivation), 4: no fauna and temperature elevated by 1.5 °C, 5: temperature elevated by 3 °C (refer to SI 2 for the derivation), 6: no fauna and temperature elevated by 3 °C) were run with the same input parameters and variables.

For the start conditions, the values per group were averaged for the first date, for OS = detritus, BOC = COD, and prokaryotes. In contrast to all the other variables, at some dates, fauna is 0 in a well at a date, even if at other dates there is fauna in the same well and group. Thus, in this case, the simulation was started artificially with the average at the first date when measured fauna biomass was above 0, because otherwise, fauna stays 0 because there is no biomass to start with that can digest etc.

With the growth parameters, a lot of gross assumptions were made which might not hold up. But we need to estimate ecosystem services and how they react to environmental changes now. Where more adequate parameters are lacking, we tried to make sure that the values were at least based on reasoning which at every step may be overhauled once more adequate data are available.

Yield, Michaelis Menten half-saturation coefficient, and maximum growth rate $\mu_{max,MO}$ ($\mu_{max}$ = maximum growth rate; MO = microorganisms) were taken for the most competitive microorganism in the comparison conducted by Schmidt et al. (2018). For simplicity and due to lack of data, we assumed these parameters would be similar for humic acids. The acetate-related values of g (acetate) were converted to mol (acetate) by dividing by the mole mass of acetate: 60 g $mol^{-1}$. The biomass stoichiometrical formula was assumed to be on average $C_1 H_{1.8} O_{0.5} N_{0.2}$ (Heijnen, 2006; see above), i.e. a molar mass of 24.6 g $mol^{-1}$, and COD of 1.07 mol COD (mol biomass) $^{-1}$ (see above).

Microbial yield when feeding on acetate was shown to be 10.6 g cell carbon per mol acetate for *Comamomonas testosteroni* (Gerritse et al., 1992). This was divided by the molar mass of carbon of 12 g $mol^{-1}$ and multiplied by 1.07 mol COD (mol biomass)$^{-1}$ (see above) and divided by 2 mol COD (mol acetate)$^{-1}$: yield = 0.23 mol COD of dry mass (mol COD acetate)$^{-1}$. Michaelis Menten half-saturation coefficient $K_{ac}$ for acetate oxidation for this organism was given as 4.3 µM; i.e. µmol $L^{-1}$, and was multiplied by $1.07*10^{-6}$ to get $0.46 *10^{-6}$ mol COD of acetate. This value derived at 30°C (Gerritse et al., 1992) was temperature-corrected as explained in S3.

We modelled that prokaryotes take up only COD and for each time step, we set up the model so that the $k_1$th fraction of detritus was broken down physically to become COD (termed "mineralization" in Soetaert & Herman 2009). This step took account of the fact that the majority of carbon sources are not degradable by microorganisms in the short term. We assumed this to be independent of temperature –updating this assumption may be one of the next steps once this proof-of-principle study has shown satisfactory results. $k_1$ was set to 0.0001 like in Soetaert & Herman (2009).

### S7: Process model

The hybrid Monod/ Verhulst models (Schlogelhofer et al., 2021) was implemented in the form

| | |
|---|---|
| maximum uptake by $MO_{het}$ [COD mol $L^{-1}$] | maximum growth rate per day at respective temperature [COD mol $L^{-1}$ $day^{-1}$] * COD of the previous day [COD mol $L^{-1}$] ( (COD of the previous day [COD mol $L^{-1}$])$^{-1}$ + $K_{ac}$ at respective temperature) * $MO_{het}$ of the previous day [COD mol $L^{-1}$] *time step [day] * (CC_group_MO_g – MO_het_ti_minus_1) CC_group_MO_g$^{-1}$ *d |

where

| | |
|---|---|
| $MO_{het}$ | heterotrophic microorganisms |
| CC_group_MO_g | carrying capacity of $MO_{het}$ per aquifer group g; assumption: the maximum observed $MO_{het}$ [COD mol $L^{-1}$] over the period of the field study, assuming that the microorganisms had maximum been growing to the carrying capacity. |
| $K_{ac}$ at respective temperature | derived from the $K_S$ that Gerritse et al. (1992) calculated for the experimental temperature of 30°C; see "S3: Derivation of the temperature influences … " |
| maximum uptake rate per day at respective temperature | derived according to Schmidt et al. (2018) from the $\mu_{max}$ that Gerritse et al. (1992) calculated for the experimental temperature of 30°C ; see "S5: Derivation of the temperature influences … " |

This uptake by microorganisms was transformed to microorganisms` biomass via the yield with which microorganisms can make use of the up-taken substrate. The yield was taken from Schmidt et al. (2018) and had the value of 0.4601 g COD of dry mass of microorganisms $g^{-1}$ COD acetate based on the experiments by Gerritse et al. (1992)(see above), assuming that this value would approach that for humic acids well enough.

The food uptake of fauna was modelled with the same hybrid Monod / Verhulst models (Schlogelhofer et al., 2021) as the microbial uptake, with the analogous variables and factors. For fauna, as well, the maximum found fauna concentrations were set as the carrying capacity.

The carbon ingestion rates measured by Di Lorenzo et al. (2025) were 0.33 and 0.73 µg C $d^{-1}$ (individual *Diacyclops belgicus* and *D. crassicaudis crassicaudis)* $^{-1}$. Assuming an average *Diacyclops* individual dry mass of 0.0005 mg (Di Lorenzo et al., 2025), i.e. 0.0000005 g $L^{-1}$, using a biomass molar mass of 24.6 g $mol^{-1}$ for the niphargid mass (see above), 1.075 mol COD $mol^{-1}$ humic acid (see above) and 1.05 mol COD $mol^{-1}$ biomass (see above), a ratio of dry to wet weight of 0.1, and further assuming COD of 1 mol C to be 1 (1 C + 1 $O_2$ → 1 $CO_2$) this translates to 0.1 and 0.3 mol COD carbon (mol COD dry mass)$^{-1}$ $d^{-1}$, respectively.

Navel et al. (2011) had measured a mean assimilation of 7.5 ± 5.9 mg OM $day^{-1}$ (g dry *Niphargus rhenorhodanensis*) $^{-1}$. Using a molar mass of 227.172 g $mol^{-1}$ for humic acid (see above), and a biomass molar mass of 24.6 g $mol^{-1}$ for the niphargid mass (see above), 1.075 mol COD $mol^{-1}$ humic acid (see above) and 1.05 mol COD $mol^{-1}$ biomass (see above) this translates to 0.0008 ± 0.0007 mol COD OM $day^{-1}$ (mol COD dry niphargid)$^{-1}$ for the Navel study. This means that there is a difference of 3 orders of magnitude between the two studies, but this difference may be due to different taxa having been used, in different study setups etc.

A third study related uptake of carbon to N content of four taxa (Mermillod-Blondin et al., 2025). It showed on average 0.15 mg of OC from biofilm per day per mg of asellid N content. Relating this to C content of fauna, although the authors stress that this is inadequate, and making similar transformations as above, yielded 0.00012 mol biofilm COD (mol fauna COD $d^{-1}$)$^{-1}$.

Taking the approximate average of 0.1 and 0.3 mol COD carbon (mol COD dry mass)$^{-1}$ $d^{-1}$ from the Di Lorenzo study, the average 0.00012 mol biofilm COD (mol fauna COD $d^{-1}$)$^{-1}$. (Mermillod-Blondin et al., 2025), and 0.0008 mol COD OM day$^{-1}$ (mol COD dry niphargid)$^{-1}$ for the Navel study, results in 0.08 mol COD (mol COD dry mass)$^{-1}$ $d^{-1}$. Multiplying this value with an average concentration of 6.9E-06 mol fauna COD dry mass $L^{-1}$ in the River Fulda plain would yield an average rate of 5.8E-07 mol COD $L^{-1}$ $d^{-1}$ taken up by fauna. Assuming that these values were representative of the temperatures in the River Fulda floodplain, and assuming that this mean uptake rate can be equated to the daily half saturation concentration, yields $K_{S,MO\_het}$ = 5.8E-07 mol COD $L^{-1}$.

"The growth rate of amphipods reported as bacterial biomass ($\mu_2$) was" Foulquier et al. (2010) used a polynomial relationship for deriving amphipod biomass growth from bacterial and existing amphipod biomasses with a growth rate with the unit 0.35 per µg C bacteria per day. This cannot be translated into a maximum growth rate within the Monod equation. Assuming that the fauna growth rate is a hundredth of the prokaryote growth rate, we set the fauna growth rate to 0.103 per day.

For the grazing of fauna on microorganisms, the temperature-dependent relationships will be different, since the optimal temperature for fauna will be lower than that of the diverse microorganisms' community where "everything is everywhere … the environment selects" (Baas Becking, 1934). Di Lorenzo et al. (2025) showed a decrease in uptake by *Diacyclops* with increasing temperatures. They studied two temperatures (14°C and 17°C) which are considerably higher than the average former (8.5°C) and prognosed (10°C; 11.5 °C) temperatures in the Fulda floodplain so that we did not build a relationship from these two points to deduce a relationship for the River Fulda floodplain.

Foulquier et al. (2010) found the yield of fauna grazing in microorganisms to be 0.06, i.e. 0.21 mol COD fauna dry mass (mol COD microorganisms)$^{-1}$. That is only about a third of the value of microorganisms taking up acetate. And it is only a fifth of the growth efficiency of 30%, i.e. 0.3 mol COD biomass (mol COD microorganisms)$^{-1}$, estimated by Ikeda & Motoda (1978) – a value that has been used repeatedly since (e.g. Bode et al., 2018; Di Lorenzo et al., 2025). However, Ikeda & Motoda (1978) studied marine systems which can be assumed to be more productive than groundwater, since they are, up to a depth of ca. 50 m, not light-limited and nutrients and carbon exchange more freely and frequently. This might lead to higher quality food than in groundwater. Using the very conservative value of 0.06 ensures that we are not over estimating fauna's role in the food web but rather underestimating it.

The fauna excretion rate factor was set to 0.001, i.e. every day, 0.1 % of the dry mass was excreted. That was, if the analogy may be drawn, a tenth of the proportion given in Soetaert & Herman (2009). These authors studied marine systems which at the sea surface can be believed to be more productive than groundwater. A factor of 0.001 is more conservative.

Fauna mortality rate was set to a low value of 1E-07 (mol COD $L^{-1}$)$^{-1}$ day$^{-1}$, i.e. a tenth of the average biomass, based on them being K strategists with a long life span (Hose et al., 2022; Saccò et al., 2024). Mortality was then calculated as mortality rate multiplied by the square of the fauna dry mass mol COD $L^{-1}$, resulting in mol COD $L^{-1}$ $day^{-1}$, which was subtracted from the respective fauna dry mass concentration in mol COD $L^{-1}$ for the respective day, in the case of daily modelling steps.

In addition, a temperature-dependent mortality rate was developed, since it is known that mortality increases up to 100%, depending on the duration, when environmental temperature is increased by as little as 4 °C (Brielmann et al., 2011; Di Lorenzo & Reboleira, 2022; Spengler, 2017; Spengler & Hahn, 2018). Brielmann et al. (2011) showed that 100% of the *Proasellus cavaticus* and *Niphargus inopinatus* test organisms had survived at 12 °C, between 50 and 100% at 16 °C, and between 0 and 50% at 20 °C. Assuming that half of the fauna population behaves like amphipods and half like asellids, this would mean 100% survival at 12 °C, 75% survival at 16 °C, and 25 % at 20 °C. With a starting point of 12 °C and an endpoint of 20 °C, a proportion of 0.25 of the population survived after an increase by 8 °C, in other words, a proportion of 0.75 died. Thus, assuming roughly a linear response, for each degree of increase, 0.75 $8^{-1}$ = 0.094, i.e. roughly 1 % of the individuals died.

In the River Fulda floodplain, groundwater temperatures were regularly lower than 12 °C. Calculating the absolute deviation of the reference temperature of 12 °C, would mean that temperatures of e.g. 7 °C would have been as lethal as temperatures of 17 °C, which is unrealistic. However, we lack reliable relationships below 12°C. At cold temperatures, fauna succumbs to cold rigor, but this is not deadly – at least not in the short run (Brielmann et al., 2011). Therefore, we implemented the relationship only for temperatures above 12 °C and set the temperature-related mortality to 0 at temperatures below 12°C. This needs to be refined in future experiments.

The growth in each time step was added to the biomass (detritus, COD, microorganisms, or fauna) of the previous time step. Whenever the resulting new biomass was negative, because the growth was more negative due to loss terms, than the biomass of the previous step, the net biomass was artificially set to a minute value of 0.00000000001 mol COD $L^{-1}$. Negative biomass is not defined, and assuming that there would always be at least a minute import or reactivation of dormant cells, we considered this step a valid approximation. Further studies must include import and export of biomass by transport.

### S8: Overview over the scenarios based on the reference scenario in the main paper, Fig. 1

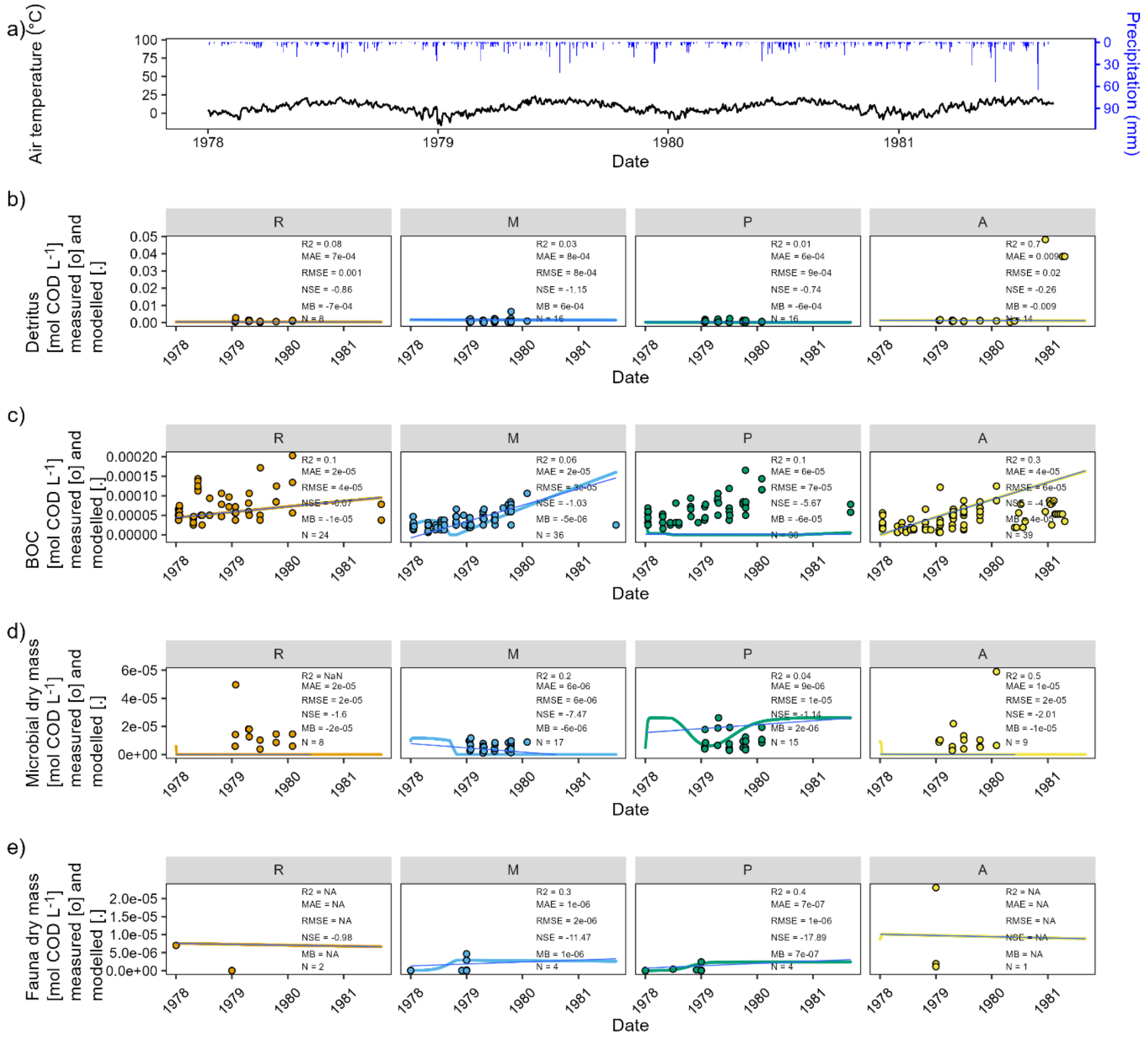


*Figure S3: Reference scenario, as in Fig. 1 in the main paper, but here with added trend line in blue. These trends lines, also from the subsequent figures, were used in Fig. 2 in the main paper. River-near = "R", mixing zone = "M", plume = "P", agricultural area = "A". a) Air temperature (black) and precipitation in Fulda Horas meteorological station 1526 (blue) for the four years covering the study period. b), c), d), e): measured (dots) and modelled (lines) carbon compounds in the four well groups (refer to text and Fig. S1). This scenario is referred to as the "Reference" scenario. Please note that the highest measured values in fauna dry mass (1.3*10$^{-04}$ in P03, group „R"; on 2 January 1979) was omitted from the plot to enable visualization of the modelled time course.*

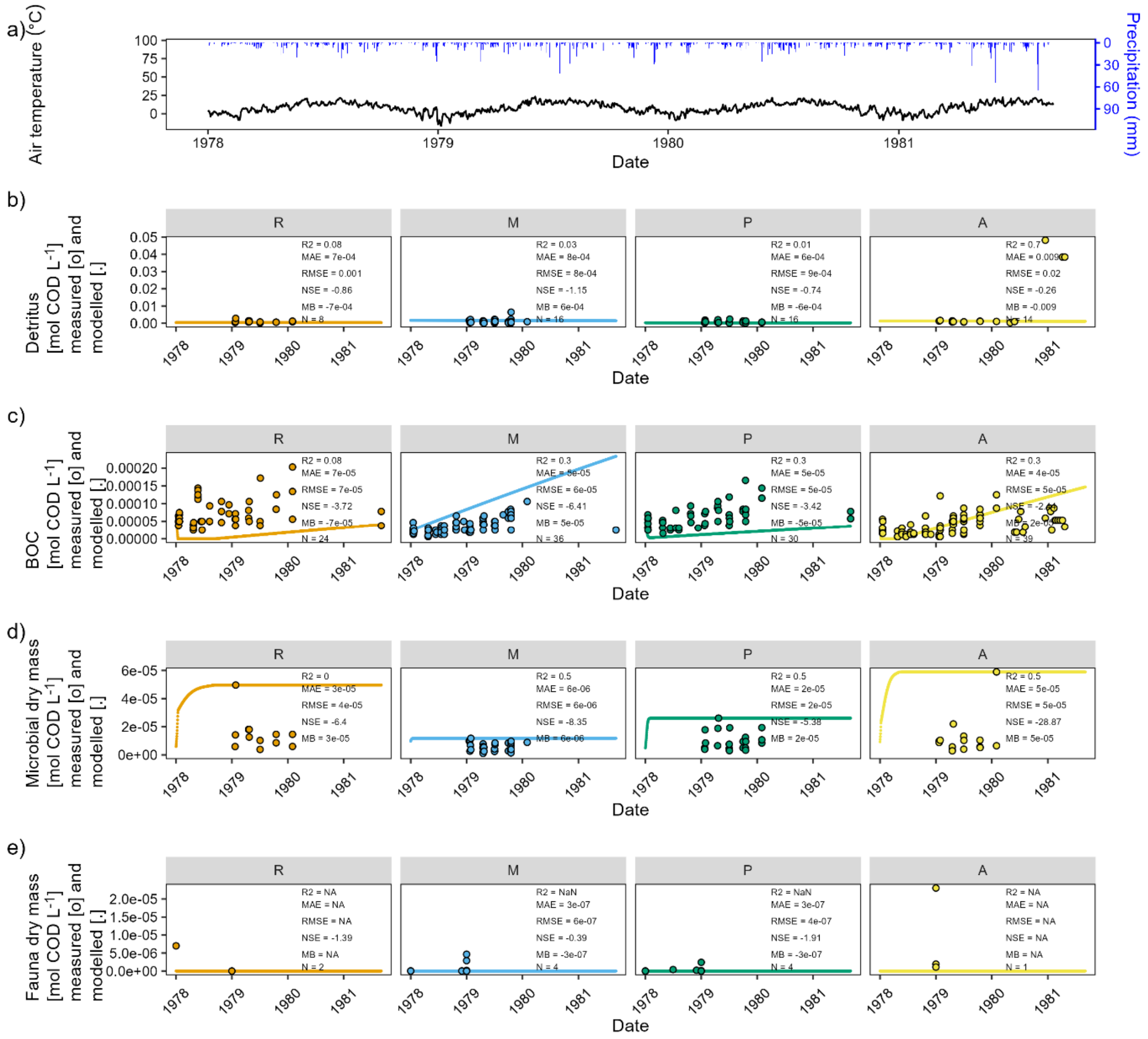


*Figure S4: Scenario "no fauna". In the model setup the results of which are shown in Fig. 1 in the main paper, fauna was set to 0 without recruitment over time. River-near = "R", mixing zone = "M", plume = "P", agricultural area = "A". a) Air temperature (black) and precipitation in Fulda Horas meteorological station 1526 (blue) for the four years covering the study period. b), c), d), e): measured (dots) and modelled (lines) carbon compounds in the four well groups (refer to text and Fig. S1). Please note that the highest measured values in fauna dry mass (1.3*10$^{-04}$ in P03, group „R"; on 2 January 1979) was omitted from the plot to enable visualization of the modelled time course.*

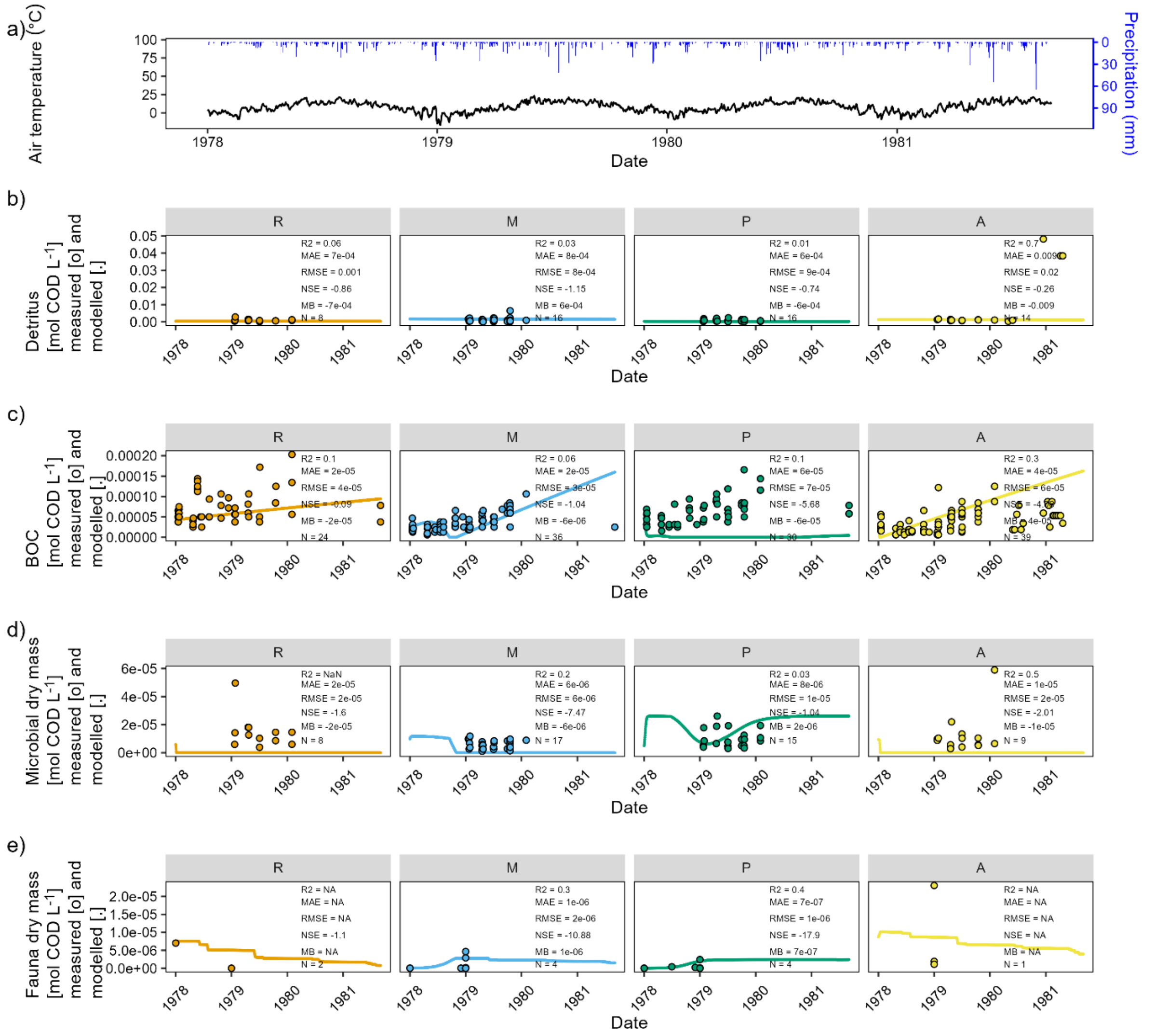


*Figure S5: Scenario "+1.5 °C". In the model setup the results of which are shown in Fig. 1 in the main paper, fauna was set to 0 without recruitment over time. River-near = "R", mixing zone = "M", plume = "P", agricultural area = "A". a) Air temperature (black) and precipitation in Fulda Horas meteorological station 1526 (blue) for the four years covering the study period. b), c), d), e): measured (dots) and modelled (lines) carbon compounds in the four well groups (refer to text and Fig. S1). Please note that the highest measured values in fauna dry mass (1.3*10$^{-04}$ in P03, group „R"; on 2 January 1979) was omitted from the plot to enable visualization of the modelled time course.*

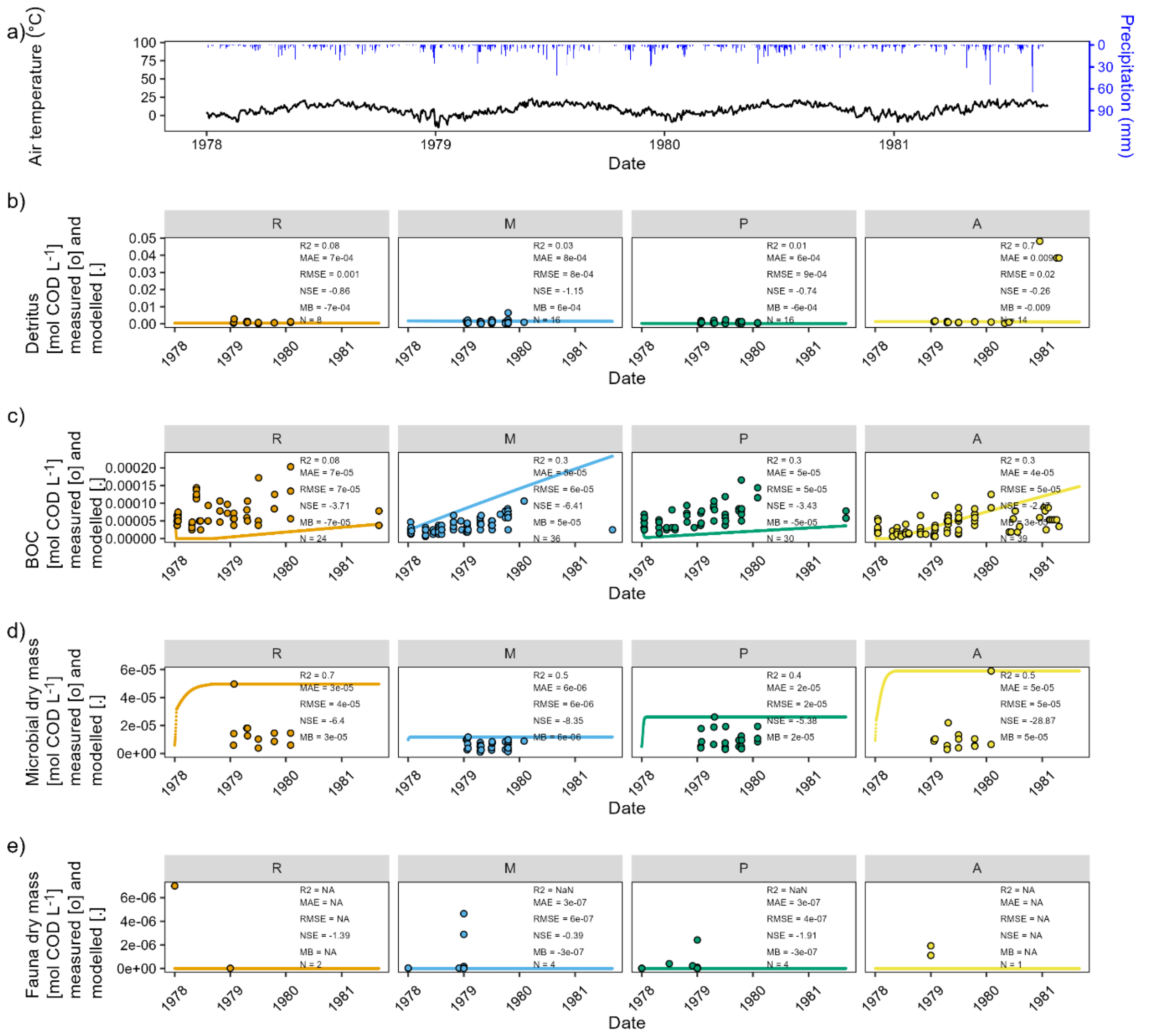


*Figure S6: Scenario "no fauna at +1.5 °C". In the model setup the results of which are shown in Fig. 1 in the main paper, fauna was set to 0 without recruitment over time. River-near = "R", mixing zone = "M", plume = "P", agricultural area = "A". a) Air temperature (black) and precipitation in Fulda Horas meteorological station 1526 (blue) for the four years covering the study period. b), c), d), e): measured (dots) and modelled (lines) carbon compounds in the four well groups (refer to text and Fig. S1). Please note that the highest measured values in fauna dry mass (1.3*10$^{-04}$ in P03, group „R"; on 2 January 1979) was omitted from the plot to enable visualization of the modelled time course.*

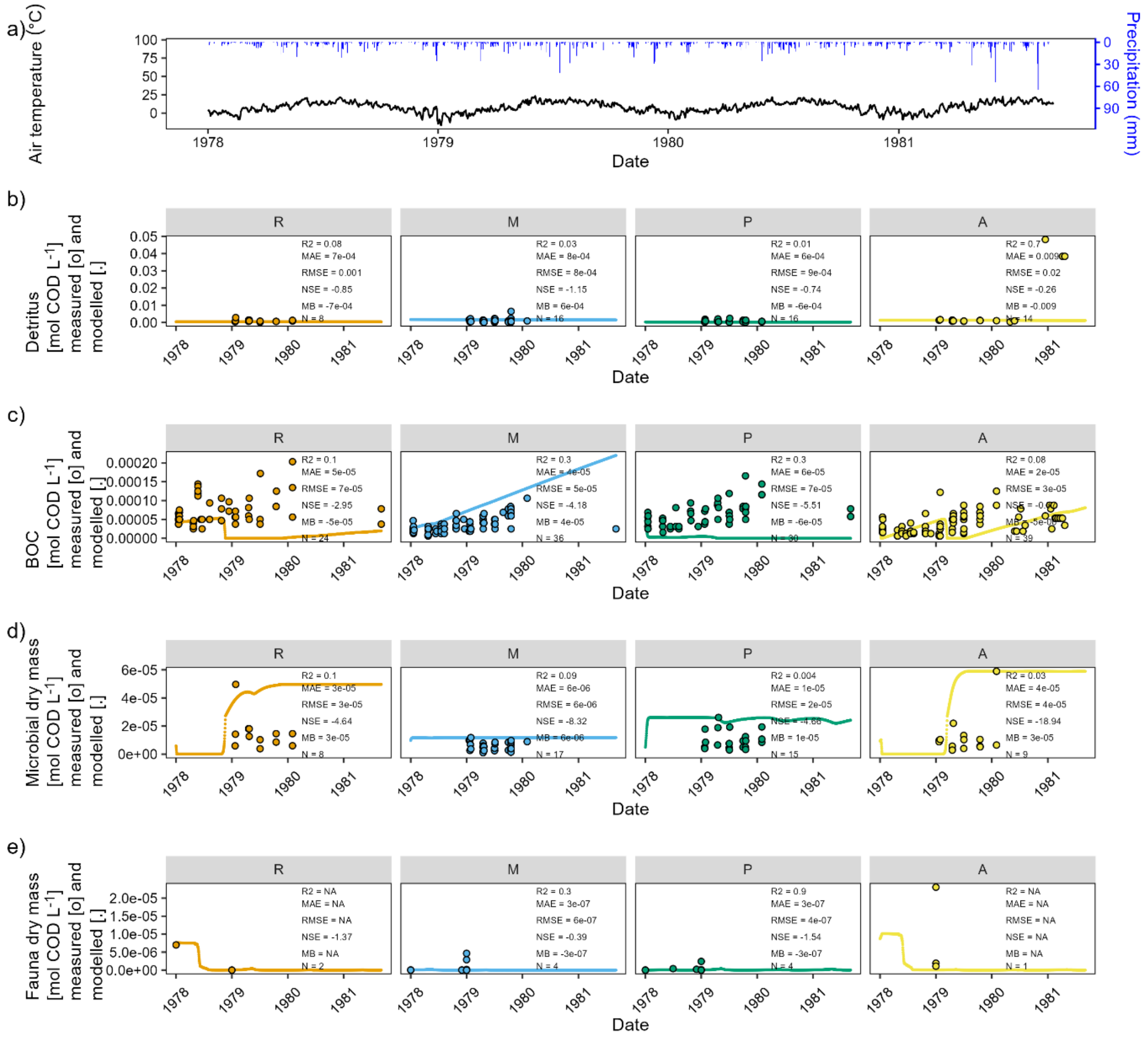


*Figure SI 2: Scenario "+3 °C". In the model setup the results of which are shown in Fig. 1 in the main paper, fauna was set to 0 without recruitment over time. River-near = "R", mixing zone = "M", plume = "P", agricultural area = "A". a) Air temperature (black) and precipitation in Fulda Horas meteorological station 1526 (blue) for the four years covering the study period. b), c), d), e): measured (dots) and modelled (lines) carbon compounds in the four well groups (refer to text and SI Fig. 1). Please note that the highest measured values in fauna dry mass (1.3*10$^{-04}$ in P03, group „R"; on 2 January 1979) was omitted from the plot to enable visualization of the modelled time course.*

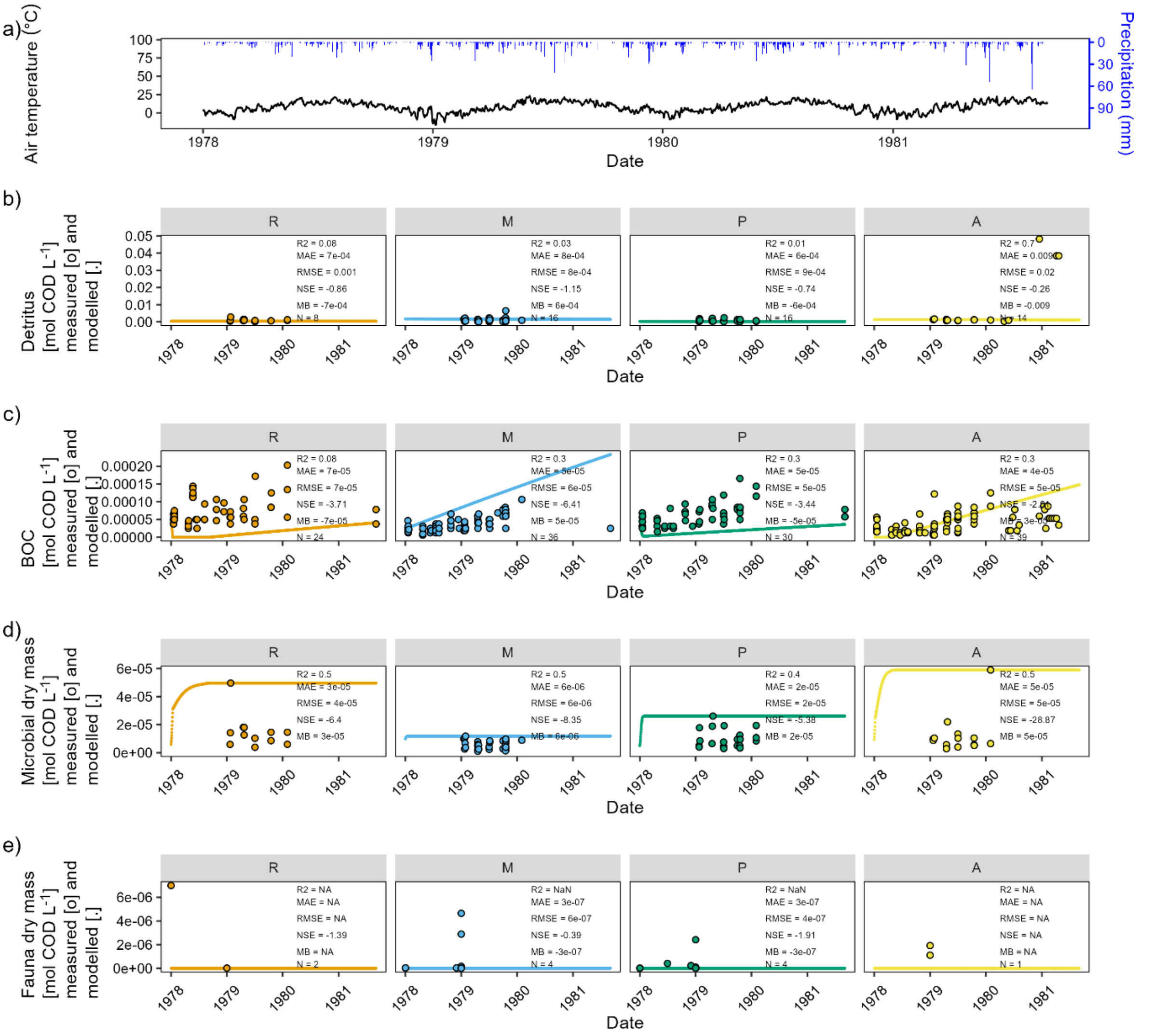


*Figure S8: Scenario "no fauna at +3 °C". In the model setup the results of which are shown in Fig. 1 in the main paper, fauna was set to 0 without recruitment over time. River-near = "R", mixing zone = "M", plume = "P", agricultural area = "A". a) Air temperature (black) and precipitation in Fulda Horas meteorological station 1526 (blue) for the four years covering the study period. b), c), d), e): measured (dots) and modelled (lines) carbon compounds in the four well groups (refer to text and Fig. S1). Please note that the highest measured values in fauna dry mass ($1.3*10^{-04}$ in P03, group „R"; on 2 January 1979) was omitted from the plot to enable visualization of the modelled time course.*

### S9: Overview over the linear trends of the modelled microbial dry mass and fauna development

Modelled microbial dry mass trends, like the BOC trends, were indistinguishable and showed the lowest slopes for the three "no fauna" scenarios (yellow and light and dark orange lines in Fig. S9). In the river-near "R" and in the agricultural "A" zones, the three fauna scenario trends were almost indistinguishable from each other. In the mixing "M" zone, the "+3°C" scenario trend was in-between that of "no fauna" and the two other "fauna" scenarios. Only in the plume "P" zone was the reference scenario that with the lowest slope of the three "fauna" scenarios (Fig. S10). In summary, microbial growth was steepest under the presence of fauna.

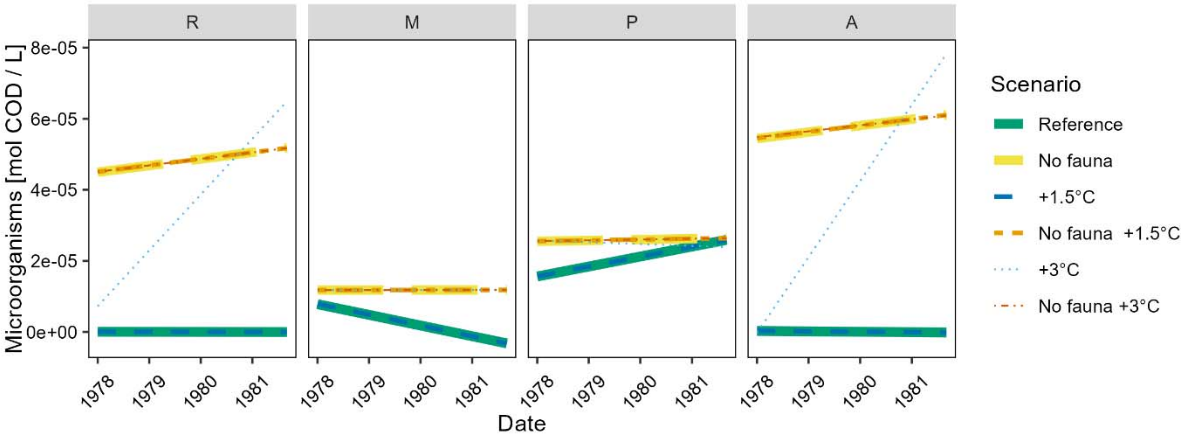


*Figure S9: Trends through the modelled microbial dry mass in the river Fulda floodplain for the six scenarios: "Reference": input variables according to Marxsen et al. (2021); "No fauna": setting all fauna variables to 0; "+1.5°C": raising all groundwater temperatures by 1.5°C; "No fauna +1.5°C": setting all fauna variables to 0 and raising all groundwater temperatures by 1.5°C; "+3°C: raising all groundwater temperatures by 3°C; "No fauna +3°C": setting all fauna variables to 0 and raising all groundwater temperatures by 3°C. The four sub plots depict the four aquifer groups: river-near = "R", mixing zone = "M", plume = "P", agricultural area = "A". Note the different scales of the y axes.*

Faunal dry mass was of course 0 in all "no fauna" scenarios and is indicated in Fig. S10 (yellow and light and dark orange lines) only for completeness. Modelled fauna dry mass trends decreased in all "fauna" scenarios. The reference scenario trend was always the one with the highest intercept and steepest decrease, and, vice versa, the "+3°C" scenario was the one that came closest to the "no fauna" scenario zero trends.

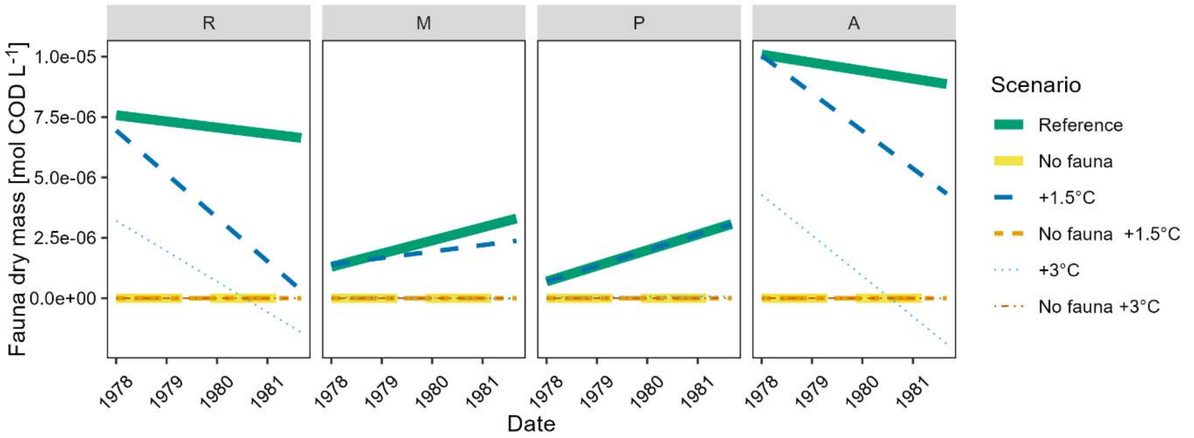


*Figure S10: Trends through the modelled fauna dry mass in the river Fulda floodplain for the six scenarios: "Reference": input variables according to Marxsen et al. (2021); "No fauna": setting all fauna variables to 0; "+1.5°C": raising all groundwater temperatures by 1.5°C; "No fauna +1.5°C": setting all fauna variables to 0 and raising all groundwater temperatures by 1.5°C; "+3°C: raising all groundwater temperatures by 3°C; "No fauna +3°C": setting all fauna variables to 0 and raising all groundwater temperatures by 3°C. The four sub plots depict the four aquifer groups: river-near = "R", mixing zone = "M", plume = "P", agricultural area = "A". Note the different scales of the y axes. The zero lines for the scenarios without fauna are only added for completion.*

### S10: Parameters of the linear fits to the scenarios

*Table S1: Linear model fits to the scenario runs. River-near = "R", mixing zone = "M", plume = "P", agricultural area = "A". BOC = biologically oxidizable carbon; MO_het = dry mass COD heterotrophic microorganisms; "conc." = concentration. For the calculations, the value of "Modelled concentration at end time point was changed from the minute value of $2.75+10^{-271}$ for the dry mass COD heterotrophic microorganisms in group P in the reference scenario to the low value of $1*10^{-11}$ that occurred in other simulations as well. All % changes and factors "by which modelled end concentration changed from reference modelled end concentration" are thus gross under estimates. The lines for BOC, for which the ecosystem service of carbon degradation is reported and discussed, are indicated in bold.*

| Scenario | Group | Vari-able | Inter-cept of fit | Slope of fit [mol COD $L^{-1}$ $d^{-1}$] | p-value of fit | Model-led conc. time 0 [mol COD $L^{-1}$] | Modelled conc. time end [mol COD $L^{-1}$] | Fitted conc. time 0 [mol COD $L^{-1}$] | Fitted conc. time end [mol COD $L^{-1}$] | Modelled conc. at end minus modelled conc. time 0 [mol COD $L^{-1}$ $d^{-1}$] | % change at modelled end com-pared to modelled conc. time 0 | Factor by which modelled end conc. changed from refe-rence modelled end conc. |
|---|---|---|---|---|---|---|---|---|---|---|---|---|
| Reference | **R** | **BOC** | **-6.8E-05** | **3.8E-08** | **0.0E+00** | **5.1E-05** | **9.5E-05** | **4.4E-05** | **9.6E-05** | **4.44E-05** | **87.4** | |
| | R | MO_het | 2.3E-07 | -5.9E-11 | 5.7E-04 | 5.9E-06 | 1.0E-11 | 5.3E-08 | -2.6E-08 | -5.94E-06 | -100.0 | |
| | R | fauna | 9.6E-06 | -7.0E-10 | 0.0E+00 | 7.0E-06 | 6.6E-06 | 7.6E-06 | 6.6E-06 | -3.59E-07 | -5.1 | |
| | **M** | **BOC** | **-3.4E-04** | **1.1E-07** | **0.0E+00** | **2.8E-05** | **1.6E-04** | **-8.0E-06** | **1.5E-04** | **1.32E-04** | **470.2** | |
| | M | MO_het | 3.2E-05 | -8.2E-09 | 4.3E-211 | 9.8E-06 | 1.0E-11 | 7.8E-06 | -3.2E-06 | -9.81E-06 | -100.0 | |
| | M | fauna | -3.1E-06 | 1.5E-09 | 1.5E-151 | 2.2E-08 | 2.6E-06 | 1.3E-06 | 3.3E-06 | 2.58E-06 | 11608.7 | |
| | **P** | **BOC** | **2.2E-06** | **-2.2E-10** | **3.8E-01** | **4.7E-05** | **5.5E-06** | **1.6E-06** | **1.3E-06** | **-4.14E-05** | **-88.3** | |
| | P | MO_het | -6.8E-06 | 7.7E-09 | 4.1E-62 | 5.0E-06 | 2.6E-05 | 1.6E-05 | 2.6E-05 | 2.11E-05 | 425.0 | |
| | P | fauna | -4.5E-06 | 1.8E-09 | 2.3E-282 | 6.9E-09 | 2.4E-06 | 6.9E-07 | 3.1E-06 | 2.40E-06 | 34657.1 | |
| | **A** | **BOC** | **-3.6E-04** | **1.2E-07** | **0.0E+00** | **2.8E-05** | **1.6E-04** | **5.7E-07** | **1.6E-04** | **1.35E-04** | **489.5** | |
| | A | MO_het | 1.2E-06 | -3.1E-10 | 1.8E-09 | 9.3E-06 | 1.0E-11 | 2.8E-07 | -1.4E-07 | -9.32E-06 | -100.0 | |
| | A | fauna | 1.3E-05 | -9.1E-10 | 0.0E+00 | 8.7E-06 | 8.9E-06 | 1.0E-05 | 8.9E-06 | 1.73E-07 | 2.0 | |
| No Fauna | **R** | **BOC** | **-1.0E-04** | **3.3E-08** | **0.0E+00** | **5.1E-05** | **4.1E-05** | **-4.9E-06** | **3.9E-05** | **-1.01E-05** | **-19.8** | **0.43** |
| | R | MO_het | 3.0E-05 | 5.1E-09 | 3.6E-69 | 5.9E-06 | 5.0E-05 | 4.5E-05 | 5.2E-05 | 4.37E-05 | 735.2 | 4962171 |
| | R | fauna | 0 | 0 | | 0 | 0 | 0 | 0 | 0 | | |
| | **M** | **BOC** | **-4.3E-04** | **1.6E-07** | **0.0E+00** | **2.8E-05** | **2.3E-04** | **2.6E-05** | **2.4E-04** | **2.05E-04** | **729.2** | **1.45** |
| | M | MO_het | 1.2E-05 | 3.9E-11 | 1.3E-07 | 9.8E-06 | 1.2E-05 | 1.2E-05 | 1.2E-05 | 2.00E-06 | 20.4 | 1180744 |
| | M | fauna | 0 | 0 | | 0 | 0 | 0 | 0 | 0 | | |
| | **P** | **BOC** | **-6.6E-05** | **2.4E-08** | **0.0E+00** | **4.7E-05** | **3.6E-05** | **3.8E-06** | **3.6E-05** | **-1.07E-05** | **-22.9** | **6.62** |
| | P | MO_het | 2.4E-05 | 6.8E-10 | 2.3E-11 | 5.0E-06 | 2.6E-05 | 2.6E-05 | 2.6E-05 | 2.11E-05 | 425.4 | 1 |
| | P | fauna | 0 | 0 | | 0 | 0 | 0 | 0 | 0 | | |
| | **A** | **BOC** | **-3.6E-04** | **1.2E-07** | **0.0E+00** | **2.8E-05** | **1.5E-04** | **-1.1E-05** | **1.5E-04** | **1.19E-04** | **432.7** | **0.90** |
| | A | MO_het | 4.0E-05 | 5.0E-09 | 1.8E-43 | 9.3E-06 | 5.9E-05 | 5.4E-05 | 6.1E-05 | 4.96E-05 | 532.3 | 5893511 |
| | A | fauna | 0 | 0 | | 0 | 0 | 0 | 0 | 0 | | |
| +1.5°C | **R** | **BOC** | **-6.9E-05** | **3.8E-08** | **0.0E+00** | **5.1E-05** | **9.4E-05** | **4.3E-05** | **9.5E-05** | **4.36E-05** | **85.9** | **0.99** |
| | R | MO_het | 2.3E-07 | -6.1E-11 | 4.5E-04 | 5.9E-06 | 2.7E-256 | 5.5E-08 | -2.7E-08 | -5.94E-06 | -100.0 | 0.00 |
| | R | fauna | 2.1E-05 | -4.9E-09 | 0.0E+00 | 7.0E-06 | 7.7E-07 | 6.9E-06 | 3.2E-07 | -6.23E-06 | -88.9 | 0.12 |
| | **M** | **BOC** | **-3.4E-04** | **1.1E-07** | **0.0E+00** | **2.8E-05** | **1.6E-04** | **-8.1E-06** | **1.5E-04** | **1.32E-04** | **468.2** | **1.00** |
| | M | MO_het | 3.2E-05 | -8.3E-09 | 1.3E-213 | 9.8E-06 | 1.7E-261 | 7.9E-06 | -3.2E-06 | -9.81E-06 | -100.0 | 0.00 |
| | M | fauna | -7.3E-07 | 7.3E-10 | 1.4E-42 | 2.2E-08 | 1.5E-06 | 1.4E-06 | 2.4E-06 | 1.47E-06 | 6620.9 | 0.57 |
| | **P** | **BOC** | **2.7E-06** | **-3.9E-10** | **1.1E-01** | **4.7E-05** | **4.5E-06** | **1.6E-06** | **1.1E-06** | **-4.23E-05** | **-90.3** | **0.83** |
| | P | MO_het | -6.5E-06 | 7.6E-09 | 4.6E-61 | 5.0E-06 | 2.6E-05 | 1.6E-05 | 2.6E-05 | 2.11E-05 | 424.9 | 1.00 |
| | P | fauna | -4.5E-06 | 1.8E-09 | 1.9E-283 | 6.9E-09 | 2.4E-06 | 6.9E-07 | 3.1E-06 | 2.39E-06 | 34566.8 | 1.00 |
| | **A** | **BOC** | **-3.6E-04** | **1.2E-07** | **0.0E+00** | **2.8E-05** | **1.6E-04** | **5.3E-07** | **1.6E-04** | **1.35E-04** | **489.5** | **1.00** |
| | A | MO_het | 1.3E-06 | -3.4E-10 | 1.5E-09 | 9.3E-06 | 1.0E-11 | 3.0E-07 | -1.5E-07 | -9.32E-06 | -100.0 | 1.00 |
| | A | fauna | 2.3E-05 | -4.3E-09 | 0.0E+00 | 8.7E-06 | 4.0E-06 | 1.0E-05 | 4.3E-06 | -4.73E-06 | -54.5 | 0.45 |
| No Fauna +1.5°C | **R** | **BOC** | **-1.0E-04** | **3.3E-08** | **0.0E+00** | **5.1E-05** | **4.1E-05** | **-4.9E-06** | **3.9E-05** | **-9.96E-06** | **-19.6** | **0.43** |
| | R | MO_het | 3.1E-05 | 5.0E-09 | 1.8E-68 | 5.9E-06 | 5.0E-05 | 4.5E-05 | 5.2E-05 | 4.37E-05 | 735.2 | 4962171 |
| | **R** | fauna | 0 | 0 | | 0 | 0 | 0 | 0 | 0 | | |
| | M | **BOC** | **-4.3E-04** | **1.6E-07** | **0.0E+00** | **2.8E-05** | **2.3E-04** | **2.6E-05** | **2.4E-04** | **2.05E-04** | **729.2** | **1.45** |
| | M | MO_het | 1.2E-05 | 3.7E-11 | 3.5E-07 | 9.8E-06 | 1.2E-05 | 1.2E-05 | 1.2E-05 | 2.00E-06 | 20.4 | 1180744 |
| | M | fauna | 0 | 0 | | 0 | 0 | 0 | 0 | 0 | | |
| | **P** | **BOC** | **-6.6E-05** | **2.4E-08** | **0.0E+00** | **4.7E-05** | **3.6E-05** | **3.7E-06** | **3.6E-05** | **-1.07E-05** | **-22.9** | **6.62** |
| | P | MO_het | 2.4E-05 | 6.4E-10 | 1.0E-10 | 5.0E-06 | 2.6E-05 | 2.6E-05 | 2.6E-05 | 2.11E-05 | 425.4 | 1 |
| | P | fauna | 0 | 0 | | 0 | 0 | 0 | 0 | 0 | | |
| | **A** | **BOC** | **-3.6E-04** | **1.2E-07** | **0.0E+00** | **2.8E-05** | **1.5E-04** | **-1.1E-05** | **1.5E-04** | **1.20E-04** | **433.9** | **0.91** |
| | A | MO_het | 4.0E-05 | 4.9E-09 | 1.7E-42 | 9.3E-06 | 5.9E-05 | 5.5E-05 | 6.1E-05 | 4.96E-05 | 532.3 | 5893511 |
| | A | fauna | 0 | 0 | | 0 | 0 | 0 | 0 | 0 | | |

(continued overleaf)

(Tab. S1 continued)

| Scenario | Group | Variable | Intercept of fit | Slope of fit [mol COD $L^{-1} d^{-1}$] | p-value of fit | Modelled conc. time 0 [mol COD $L^{-1}$] | Modelled conc. time end [mol COD $L^{-1}$] | Fitted conc. time 0 [mol COD $L^{-1}$] | Fitted conc. time end [mol COD $L^{-1}$] | Modelled conc. at end minus modelled conc. time 0 [mol COD $L^{-1} d^{-1}$] | % change at modelled end compared to modelled conc. time 0 | Factor by which modelled end conc. changed from reference modelled end conc. |
|---|---|---|---|---|---|---|---|---|---|---|---|---|
| +3°C | R | BOC | 1.0E-04 | -2.4E-08 | 2.1E-84 | 5.1E-05 | 2.0E-05 | 3.2E-05 | 2.4E-07 | -3.08E-05 | -60.7 | 0.21 |
| | R | MO_het | -1.2E-04 | 4.3E-08 | 1.30927396 | 5.9E-06 | 5.0E-05 | 7.2E-06 | 6.5E-05 | 4.37E-05 | 735.2 | 4962167 |
| | R | fauna | 1.3E-05 | -3.5E-09 | 2.0E-118 | 7.0E-06 | 3.0E-11 | 3.2E-06 | -1.4E-06 | -7.00E-06 | -100.0 | 0.00 |
| | M | BOC | -4.2E-04 | 1.5E-07 | 0.0E+00 | 2.8E-05 | 2.2E-04 | 1.8E-05 | 2.2E-04 | 1.92E-04 | 681.6 | 1.37 |
| | M | MO_het | 1.1E-05 | 9.6E-11 | 1.5E-36 | 9.8E-06 | 1.2E-05 | 1.2E-05 | 1.2E-05 | 2.00E-06 | 20.4 | 1180744 |
| | M | fauna | 2.0E-07 | -5.1E-11 | 6.1E-89 | 2.2E-08 | 1.8E-15 | 4.9E-08 | -2.0E-08 | -2.22E-08 | -100.0 | 0.00 |
| | P | BOC | 1.7E-05 | -4.4E-09 | 1.7E-85 | 4.7E-05 | 0.0E+00 | 4.4E-06 | -1.5E-06 | -4.69E-05 | -100.0 | 0.00 |
| | P | MO_het | 3.0E-05 | -1.4E-09 | 1.8E-31 | 5.0E-06 | 2.4E-05 | 2.6E-05 | 2.4E-05 | 1.92E-05 | 386.3 | 1 |
| | P | fauna | -3.6E-08 | 2.7E-11 | 4.9E-07 | 6.9E-09 | 2.1E-09 | 4.4E-08 | 8.1E-08 | -4.85E-09 | -70.2 | 0.00 |
| | A | BOC | -1.3E-04 | 4.5E-08 | 1.3E-233 | 2.8E-05 | 8.1E-05 | 3.2E-06 | 6.4E-05 | 5.34E-05 | 193.4 | 0.50 |
| | A | MO_het | -1.7E-04 | 5.9E-08 | 0.0E+00 | 9.3E-06 | 5.9E-05 | -2.7E-07 | 7.8E-05 | 4.96E-05 | 532.3 | 5893445 |
| | A | fauna | 1.8E-05 | -4.6E-09 | 8.7E-119 | 8.7E-06 | 5.5E-10 | 4.3E-06 | -1.9E-06 | -8.69E-06 | -100.0 | 0.00 |
| No Fauna +3°C | R | BOC | -1.0E-04 | 3.3E-08 | 0.0E+00 | 5.1E-05 | 4.1E-05 | -5.0E-06 | 3.9E-05 | -9.95E-06 | -19.6 | 0.43 |
| | R | MO_het | 3.0E-05 | 5.0E-09 | 1.3E-69 | 5.9E-06 | 5.0E-05 | 4.5E-05 | 5.2E-05 | 4.37E-05 | 735.2 | 4962171 |
| | R | fauna | 0 | 0 | | 0 | 0 | 0 | 0 | 0 | | |
| | M | BOC | -4.3E-04 | 1.6E-07 | 0.0E+00 | 2.8E-05 | 2.3E-04 | 2.6E-05 | 2.4E-04 | 2.05E-04 | 729.2 | 1.45 |
| | M | MO_het | 1.2E-05 | 3.5E-11 | 9.0E-07 | 9.8E-06 | 1.2E-05 | 1.2E-05 | 1.2E-05 | 2.00E-06 | 20.4 | 1180744 |
| | M | fauna | 0 | 0 | | 0 | 0 | 0 | 0 | 0 | | |
| | P | BOC | -6.7E-05 | 2.4E-08 | 0.0E+00 | 4.7E-05 | 3.6E-05 | 3.6E-06 | 3.6E-05 | -1.07E-05 | -22.9 | 6.62 |
| | P | MO_het | 2.4E-05 | 6.0E-10 | 4.7E-10 | 5.0E-06 | 2.6E-05 | 2.6E-05 | 2.6E-05 | 2.11E-05 | 425.4 | 1 |
| | P | fauna | 0 | 0 | | 0 | 0 | 0 | 0 | 0 | | |
| | A | BOC | -3.6E-04 | 1.2E-07 | 0.0E+00 | 2.8E-05 | 1.5E-04 | -1.1E-05 | 1.5E-04 | 1.20E-04 | 435.9 | 0.91 |
| | A | MO_het | 4.1E-05 | 4.6E-09 | 1.4E-40 | 9.3E-06 | 5.9E-05 | 5.5E-05 | 6.1E-05 | 4.96E-05 | 532.3 | 5893511 |
| | A | fauna | 0 | 0 | | 0 | 0 | 0 | 0 | 0 | | |

### S11: Regional sensitivity analysis

The regional sensitivity analysis was executed separately for the four groups (Fig.s S11 – S14).

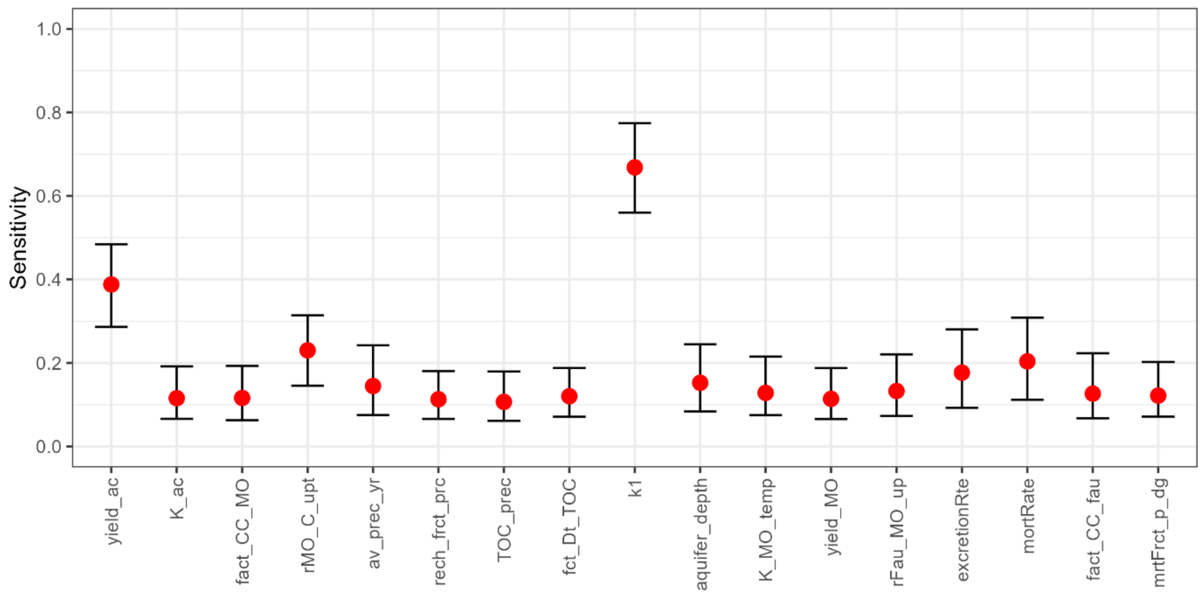


*Figure S11: Sensitivity analysis in group "M" (mixing zone). Median (red dot) and the 95% confidence interval (whiskers) of the mvd, i.e. measure-valued differentiation of the Monte Carlo boot-strapped sensitivities for the 18 parameters. Parameter acronyms are explained in Table S2.*

*Table S2: Explanation for the parameter acronyms used in Fig.s S11, S12, S13, S14.*

| Abbreviation | Full explanation |
|---|---|
| yield_ac | 4.3 microM; i.e. mol / L, i.e. mol acetate p. 1468 in Gerritse et al. (1992), i.e. multiply with 1.07 to get mol COD; will be reculculated from 30 degree from Gerritse to present temperature by factor as in Schmidt et al. (2018) |
| K_ac | 10.6 g / mol, i.e. 10.6/12 mol biomass C / mol acetate; Gerritse et al. (1992): "Expressed as grams of cell carbon produced per mole of carbon substrate consumed." This was multiplied with 1.07 mol COD / mol biomass and divided by 2 mol COD / mol acetate |
| fact_CC_MO | the factor with which maximum measured biomass of microorganisms was multiplied to derive carrying capacity of microorganisms |
| rMO_C_upt | 64 g COD / 60 g acetate, i.e. 1.067 g COD / g acetate. 2 mol COD / mol acetate. |
| mic_loss | microbial loss factor in scenarios where there is no fauna - i.e. cell respiration, grazing through the microbial loop, cell death etc. |
| av_prec_yr | average precipitation (mm / year). Plesne precipitation, to deal with Kopacek TOC in precipitation. The mean of 1402 and 1437, which are the yearly precipitaiton for Plesne and Certovo respectively (Kopacek et al., 2009: Canopy leaching .. ) |
| rech_frct_prc | the fraction of precipitation recharged to groundwater. Here a very simplified estimate which is not congruent with field measurements and has to be updated in future runs if it proves sensitive. "The infiltration rate in Chinese loess aquifers ranges between 0.19 and 0.34 m/year, representing 6–13% of the annual precipitation." Liu et al. 2024; ca. 8% Seidenfaden et al. 2023 |
| TOC_prec | concentration of total organic carbon (TOC) in precipitation: 208 mmol / m^2 / yr in Plesne, 199 mmol / m^2 / yr in Certovo; i.e. 203.5 on average; J. Kopacek et al. 2008; Kopacek, J., Turek, J., Hejzlar, J., Santruckova, H., 2009. Canopy leaching of nutrients and metals in a mountain spruce forest. Atmospheric Environment 43, 5443–5453. https://doi.org/10.1016/j.atmosenv.2009.07.031 |
| fct_Dt_TOC | the factor how many times of TOC there is carbon bound in detritus; assuming that through soil passage, the TOC from precipitation is degraded, but a lot more detritus is mobilised |
| k1 | fraction of detritus becoming BOC |
| aquifer_depth | depth of gw aquifer, in order to derive volume to which to relate input of moles |
| K_MO_temp | rate constant for bacterial uptake and oxidation of acetate Gerritse et al. (1992), assuming COD can be degraded like acetate, which is an error-prone assumption |
| yield_MO | for now for lack of better values, the same as microbes. Alternatively , one could set 0.1 as in Lindeman, but thats less.This is dry mass. |
| rFau_MO_up | this is a rough estimate and therefore is not real-temperature-adjusted at this stage |
| excretionRte | excretion rate of fauna; constant rate; excretion is biodegradable organic carbon (BOC) and added to that pool |
| mortRate | mortality rate of fauna; constant rate; freshly dead fauna is added to the detritus pool |
| fact_CC_fau | the factor with which maximum measured biomass of fauna was multiplied to derive carrying capacity of fauna |
| mrtFrct_p_dg | 100% at 12 degrees, between 50 and 100% at 16 degrees, between 0 and 50% at 20 degrees (Brielmann 2011). Assuming that half of the fauna population behaves like amphipods and half like asellids: 100% survival at 12 degrees, (75% survival at 16 degrees,) 25 % at 20 degrees. Taking the endpoint 20 degrees, 0.25 of the population survives after an increase by 8 degrees. This means that 0.75 have died. Thus, for each degree of increase, 0.75/8 = 0.09375, roughly 1 % of the individuals die; above or below 10 degree |

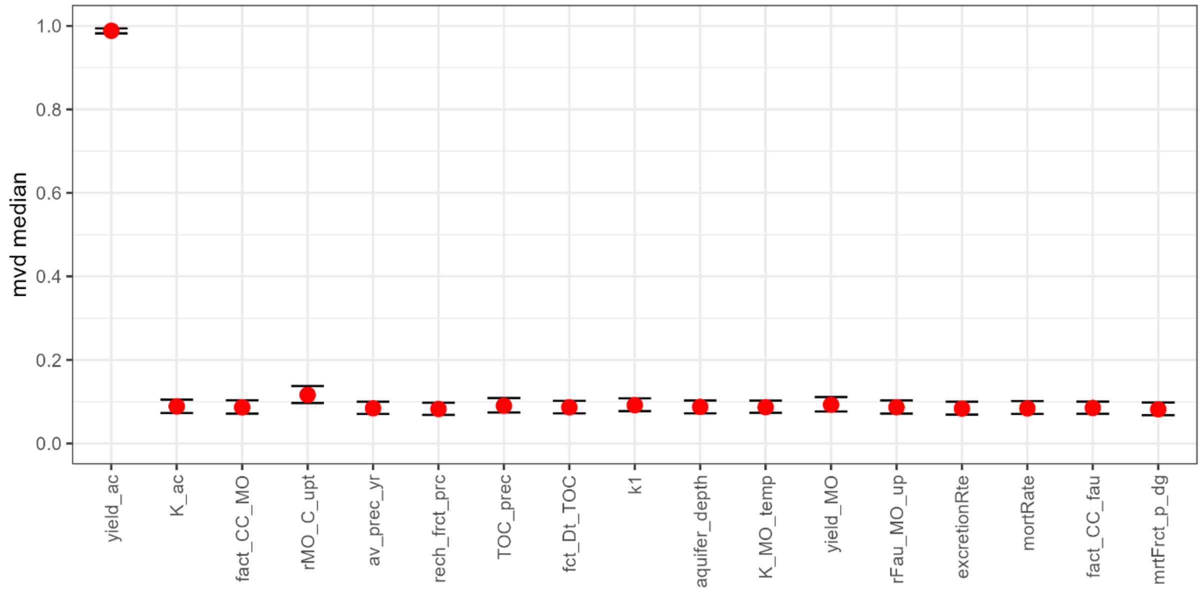


*Figure S12: Sensitivity analysis in group "P" (plume zone). Median (red dot) and the 95% confidence interval (whiskers) of the mvd, i.e. measure-valued differentiation of the Monte Carlo boot-strapped sensitivities for the 17 parameters.*

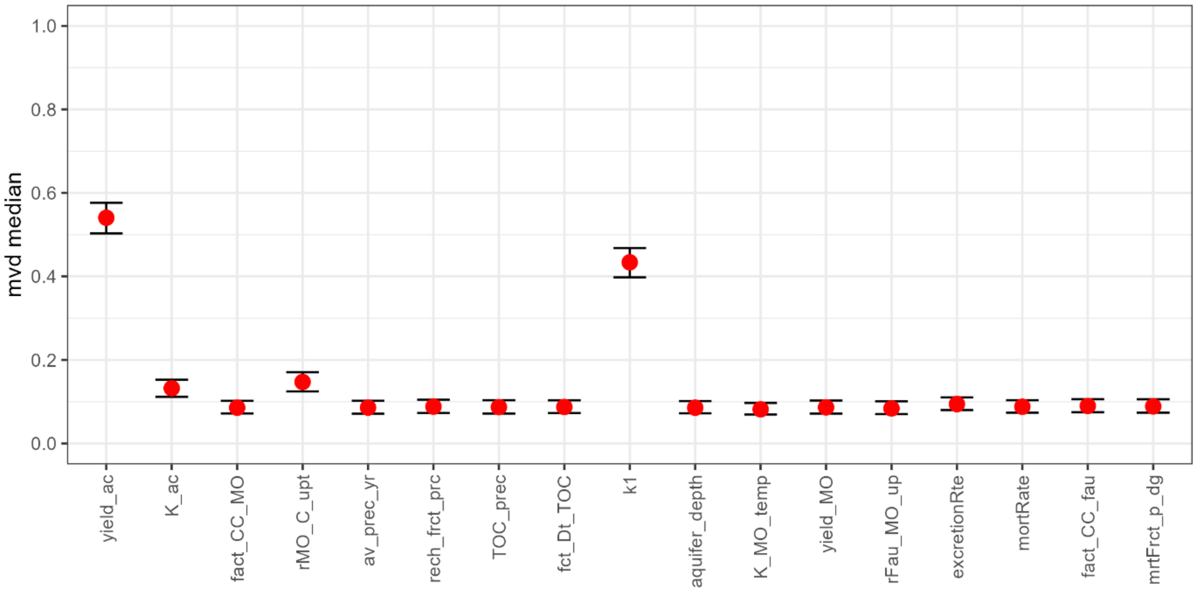


*Figure S13: Sensitivity analysis in group "R" (river-near). Median (red dot) and the 95% confidence interval (whiskers) of the mvd, i.e. measure-valued differentiation of the Monte Carlo boot-strapped sensitivities for the 18 parameters.*

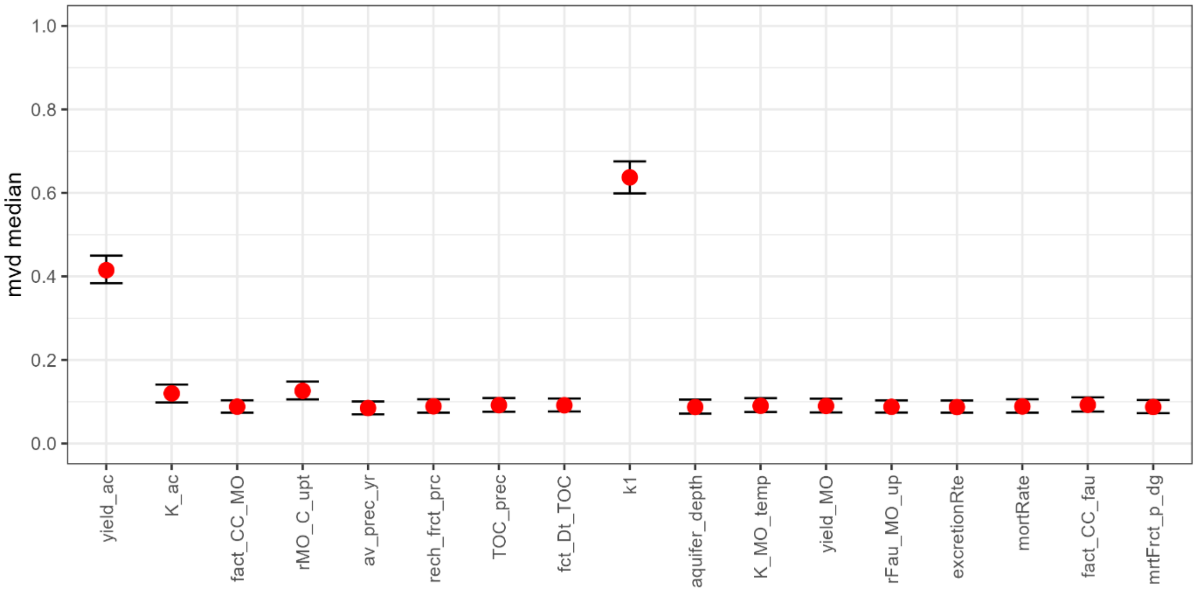


*Figure S14: Sensitivity analysis in group "A" (agriculturally influenced). Median (red dot) and the 95% confidence interval (whiskers) of the mvd, i.e. measure-valued differentiation of the Monte Carlo boot-strapped sensitivities for the 18 parameters.*

Microbial yield on carbon was the most important or second most important in all four groups. k1 was the most important or second most important only in three groups: "A", "R", and "M". Since microbial yield proved so sensitive, we ran the reference scenario (recent temperature, fauna present) with the minimum microbial yield tested in the sensitivity analysis (0.0001; Fig. S15), and the maximum microbial yield tested (0.8; Fig. S16).

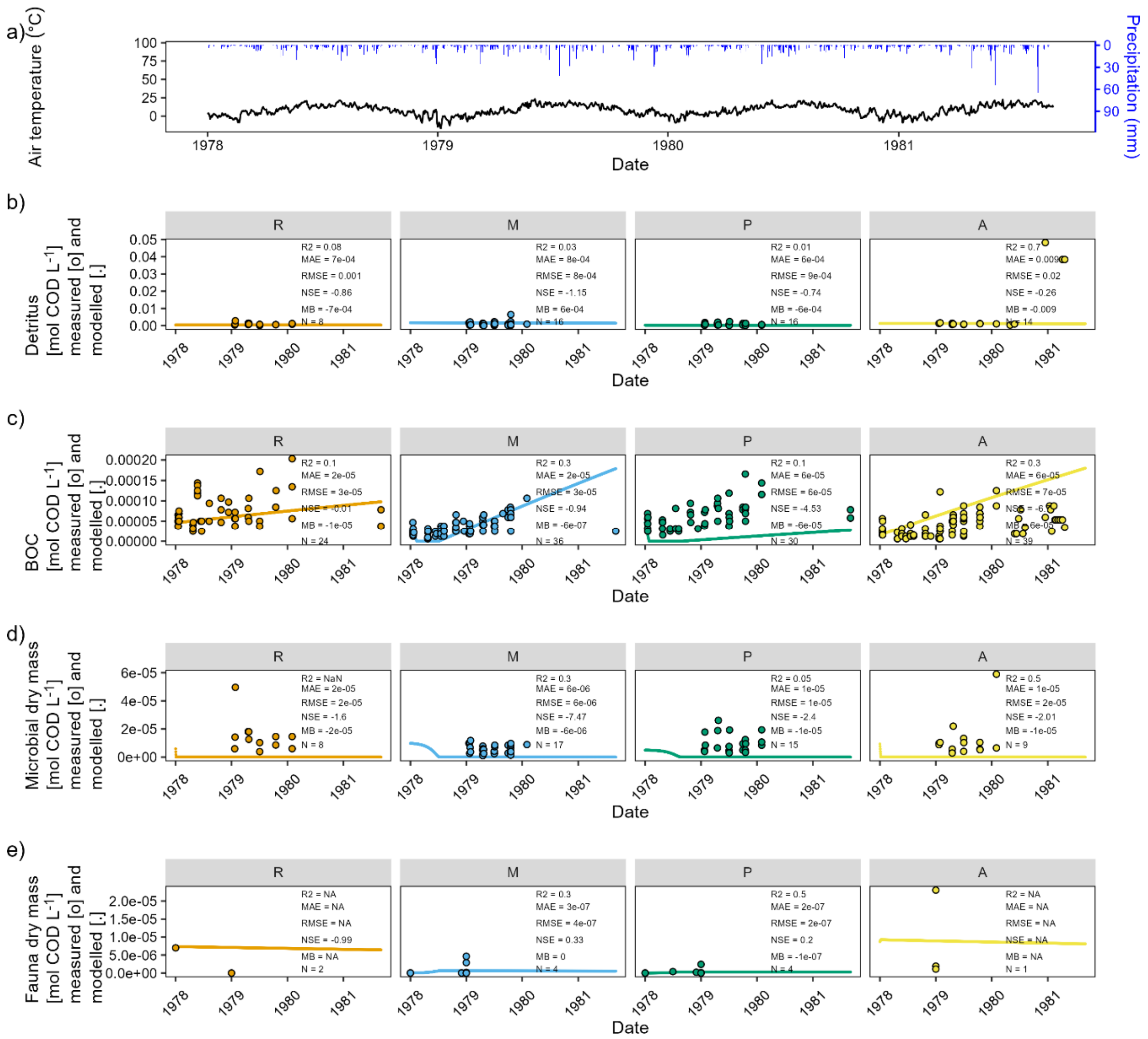


*Figure S15: Scenario based on the reference scenario, as in Figure 1 in the main paper, but here with the minimum value for microbial yield (0.0001), tested in the sensitivity analysis. River-near = "R", mixing zone = "M", plume = "P", agricultural area = "A". a) Air temperature (black) and precipitation in Fulda Horas meteorological station 1526 (blue) for the four years covering the study period. b), c), d), e): measured (dots) and modelled (lines) carbon compounds in the four well groups (refer to text and Figure S1). Please note that the highest measured values in fauna dry mass (1.3*10$^{-04}$ in P03, group „R"; on 2 January 1979) was omitted from the plot to enable visualization of the modelled time course.*

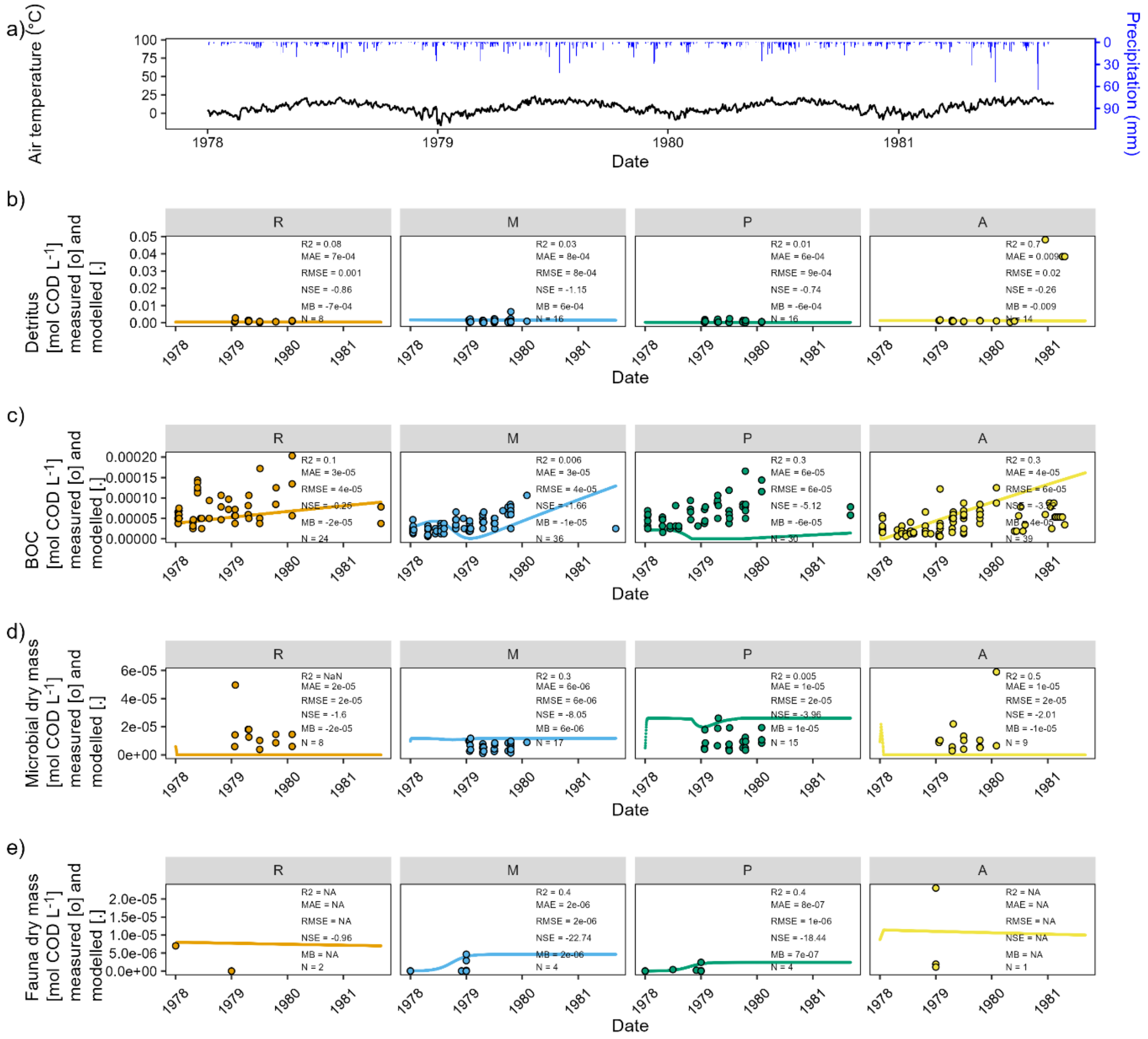


*Figure S16: Scenario based on the reference scenario, as in Fig. 1 in the main paper, but here with the maximum value for microbial yield (0.8), tested in the sensitivity analysis. River-near = "R", mixing zone = "M", plume = "P", agricultural area = "A". a) Air temperature (black) and precipitation in Fulda Horas meteorological station 1526 (blue) for the four years covering the study period. b), c), d), e): measured (dots) and modelled (lines) carbon compounds in the four well groups (refer to text and Fig. S1). Please note that the highest measured values in fauna dry mass (1.3*10$^{-04}$ in P03, group „R"; on 2 January 1979) was omitted from the plot to enable visualization of the modelled time course.*
